# snpAD: An ancient DNA genotype caller

**DOI:** 10.1101/288258

**Authors:** Kay Prüfer

**Affiliations:** Max Planck Institute for Evolutionary Anthropology, 04103 Leipzig, Germany

## Abstract

**Motivation:** The study of ancient genomes can elucidate the evolutionary past. However, analyses are complicated by base-modifications in ancient DNA molecules that result in errors in DNA sequences. These errors are particularly common near the ends of sequences and pose a challenge for genotype calling.

**Results:** I describe an iterative method that estimates genotype frequencies and errors along sequences to allow for accurate genotype calling from ancient sequences. The implementation of this method, called snpAD, performs well on high-coverage ancient data, as shown by simulations and by subsampling the data of a high-coverage Neandertal genome. Although estimates for low-coverage genomes are less accurate, I am able to derive approximate estimates of heterozygosity from several low-coverage Neandertals. These estimates show that low heterozygosity, compared to modern humans, was common among Neandertals.

**Availability:** The C++ code of snpAD is freely available at http://bioinf.eva.mpg.de/snpAD/

**Supplementary information:** Supplementary data are available at *Bioinformatics* online.

## 1 Introduction

Ancient DNA has been used to study the genomes of modern and archaic humans, mammals and plants, and has led to insights into the evolutionary past (Sarkissian *et al.*, 2015). Often, the analyses are based on sparse data. However, in some instances enough DNA molecules are preserved in ancient samples to allow for sequencing to deep coverage. This was, for instance, the case for several modern human remains (Slatkin and Racimo, 2016) and for two Neandertals and a Denisovan, extinct sister groups to present-day humans, for which genomes of at least 30-fold coverage could be generated (Meyer *et al.*, 2012; Prüfer *et al.*, 2014, 2017).

The analysis of ancient DNA is complicated by cytosine deamination, a common type of miscoding lesion accumulating in ancient DNA with increasing age and temperature (Sawyer *et al.*, 2012; Frederico *et al.*, 1990). These lesions are more frequent at the ends of ancient DNA fragments and cause cytosines to be misread as thymines (Briggs *et al.*, 2007). Laboratory methods exist that can remove this type of damage during library preparation (Briggs *et al.*, 2010). However, these methods also reduce the number of molecules that are made accessible to sequencing and are therefore best avoided when material is scarce and a high coverage genome is the aim.

The calling of diploid genotypes provides a computational means to reduce the impact of ancient DNA damage and several approaches have been published that take the characteristics of ancient DNA damage into account for calling genotypes (Jónsson *et al.*, 2013; Lindgreen *et al.*, 2014; Link *et al.*, 2017; Zhou *et al.*, 2017; Prüfer *et al.*, 2017). These approaches have fixed error rates or estimate error rates by noting differences in sequences at conserved sites or by comparing sequences to a closely related genome. The error rates are then used for quality score recalibration or directly to estimate genotype frequencies for calling genotypes.

Here, I present a different approach that jointly estimates error rates and genotype frequencies from high-coverage ancient data. Using both simulated and real ancient DNA data I demonstrate that the method is effective in dealing with high error rates in ancient DNA.

## 2 Methods

### 2.1 Implementation

SnpAD implements an iterative method that jointly estimates the frequency ofsequencing errors andthe frequency of genotypes (Fig. 1). The algorithm proceeds by first estimating by maximum likelihood the frequency of genotypes. These estimates are then used to call temporary genotypes. By comparing all sequences to these temporary genotypes, error rates are re-estimated. The three steps are iterated until the likelihood for the first step does not increase significantly. The resulting error rates and genotype frequencies can then be used to infer the most likely genotype at each position.

**Fig. 1.**
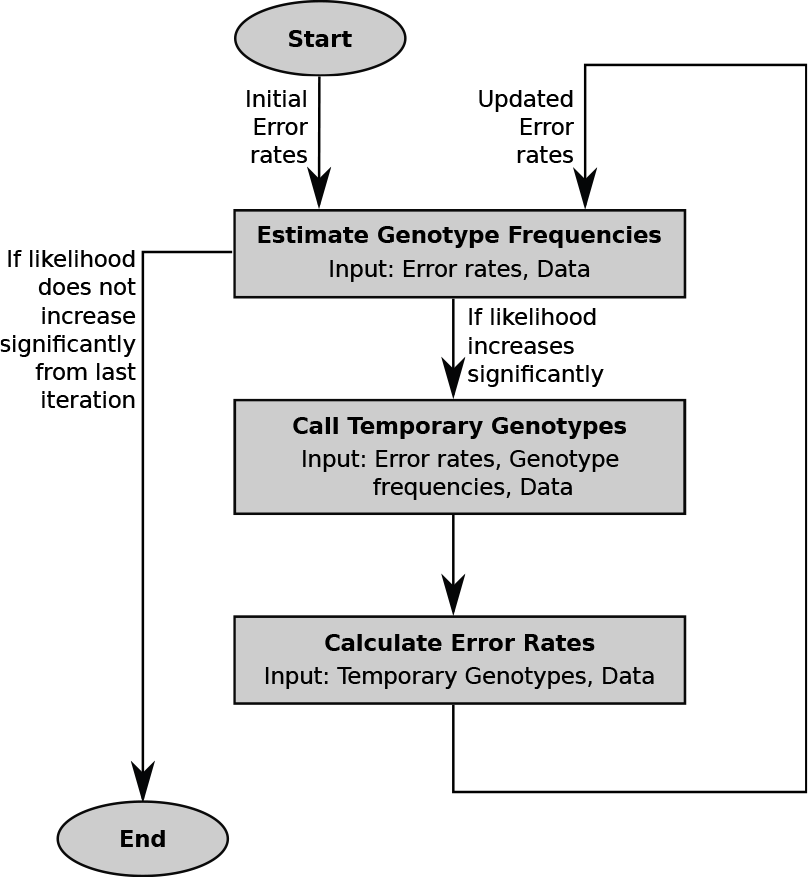
Schematic overview of the method implemented in snpAD.

### 2.1.1 Error model

The error model in snpAD assumes that the bases in the ancient sequences fall in classes that are known *a priori*, and that each class is characterized by a base substitution matrix that records the probabilities for all possible combinations of true and observed bases. In its current implementation, each base is classified by its position in the sequenced DNA fragments and optionally according to the type of sequencing library. By default, the software considers one matrix for each of the first 15 bases and last 15 bases in sequences and one matrix for bases in the interior of sequences (see Suppl. Fig. 1). However, the algorithm is independent of the classification scheme and further features of the data can be incorporated in the future.

I note that this model does not differentiate between sequencing error and ancient DNA damage, and that the probabilities given by the base quality scores are not taken into account. This choice is motivated by the fact that most ancient DNA fragments yield, due to their short length, overlapping mate pair sequences that can be merged after sequencing. The merged sequences show rarely low quality bases, indicated by a low quality score, thus rendering quality scores often largely uninformative (see Suppl. Figs. 2-3; Prüfer *et al.* 2017).

From here on, I will refer to the probability of observing base b when the true base is *B* as *P*(*b*|*B*). This probability is dependent on the strandedness of the sequence in which *b* resides and the position in this sequence. However, for ease of notation these details are omitted.

### 2.1.2 Estimating the frequency of genotypes

SnpAD estimates the frequency of all 10 possible diploid genotypes, denoted *P*(*AA*), *P*(*AC*), …, *P*(*TT*), from the data. This step assumes that the probabilities for errors are known and that sites are independent.

Genotype frequencies differ substantially: the overwhelming majority of sites are generally homozygous for one of the four bases and few sites are heterozygous. I make use of this difference by estimating the frequency of homozygous genotypes from the base composition of the data.

To estimate base composition, snpAD estimates at each position the base that likely gave rise to the observed bases given the error model. With *b*_1_,…,*b_n_* bases in *n* sequences covering a site, the score 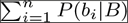 is calculated for each base *B* ∈ {*A*, *C*, *G*, *T*}. The base composition of the data, *P*(*A*), *P*(*C*), *P*(*G*), *P*(*T*), is given by the frequencies of the highest scoring bases at all sites. With the frequency of heterozygous sites *P_het_* = *P*(*AC*)+*P*(*AG*)+*P*(*AT*)+*P*(*CG*)+*P*(*CT*)+*P*(*GT*), the frequency of homozygous genotypes can be estimated using the equation

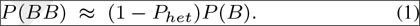

The frequency of heterozygous genotypes in the data are estimated by maximum likelihood following approaches described previously (Nielsen *et al*., 2011). Using the same notation as before, the software calculates the probability for observing *n* bases *β* = *b*_1_, …, *b_n_* when the true genotype is *B*_1_*B*_2_ as

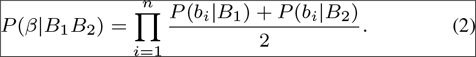

Assuming independence between sites, and considering all possible genotypes *GT* = {*AA*, *AC*, *AG*, *AT*, *CC*, *CG*, *CT*, *GG*, *GT*, *TT*}, the likelihood function is

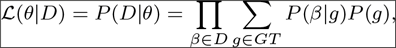

where *D* denotes the set of all sites and *θ* denotes the error rates and the genotype frequencies. The likelihood is maximized using the BOBYQA algorithm (Powell, 2009) as implemented in the library *nlopt* (Johnson, 2014) with the six heterozygous genotypes as free parameters and homozygous genotypes calculated as detailed in equation (1).

### 2.1.3 Reference bias

Both experimental and computational procedures may introduce a bias in the observed data due to the short length of ancient DNA fragments. Methods that select DNA fragments by hybridization to a DNA probe may preferentially capture DNA molecules that are identical to the probe sequence. On the other hand, since only a limited number of mismatching bases are allowed in sequence alignment, and ancient DNA sequences contain a larger proportion of mismatches due to miscoding lesions, aligning sequences may be biased towards matching the reference sequence (Prüfer *et al.*, 2010). Note that modern human contamination in archaic human genomes also contributes to an overrepresenation of reference alleles.

SnpAD offers an option to take reference bias into account when estimating genotype probabilities and when producing genotype calls. For this, a new parameter *r* (with 0.5 ≤ *r* ≤ 1) is introduced to represent the frequency at which sequences are sampled from the reference allele as opposed to the alternative alleles at heterozygous sites. For genotypes with a reference base *B*_1_ and an alternative *B*_2_ equation (2) changes to

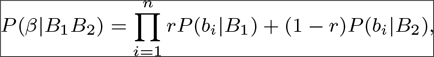

and *r* is treated as an additional free parameter during the optimization step.

### 2.1.4 Genotype calling

With the frequencies of genotypes as priors, the posterior probability for each genotype *G* at a site *β* can be calculated as

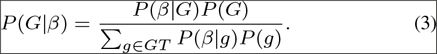

The most likely genotype is reported at each site, together with a genotype quality calculated as the Phred-scaled (Ewing and Green, 1998) log-likelihood difference between best and second best genotype.

### 2.1.5 Estimating the error model

SnpAD uses a temporary genotype call to re-estimate the error model. For this purpose, at each site the posterior probabilities for all genotypes are determined using Eq. (3).

If a site is homozygous and the genotype is known with absolute certainty, then the presence of an error could be determined by comparing the bases in each sequence to the genotype. However, genotypes are estimated using these bases so that the two are not independent. Here, I aim for an approximate solution that works despite the presence of heterozygous sites, the uncertainty in genotype calls, and the dependence of the genotype call on the underlying bases.

Sequences at heterozygous sites sample from both alleles. If each of the two alleles were to be considered the true base with equal probability, then half of the sequences would be counted as potential errors at heterozygous sites. To avoid this issue, bases that match at least one of the two alleles are not considered an error by snpAD. To solve the second issue, the uncertainty of genotype calls, snpAD counts errors proportionally to the posterior probabilities of all possible genotypes. The third issue, the dependence of a genotype call on the underlying bases, could be solved by a strategy in which each single base, in turn, is left out of the genotype call and then considered for error estimation. However, this approach would be computationally expensive. Instead, I determine using simulations which coverage is sufficiently high so that this dependence does not lead to significant bias.

## 2.2 Simulated Data

Simulated datasets with sequence coverage between 3 and 30-fold were generated to evaluate the performance of snpAD. Each dataset consists of 10 million indepdently generated sites with a fixed sequence coverage per site. A site was chosen to be heterozygous with probability 8 × 10^−4^ (2 × 10^−4^ for transitions, *CT* and *GA*, and 1 × 10^−4^ for transversions). Genotypes *CC*, *GG* had a frequency of 0.1998 and *AA*, *TT* of 0.2998.

Bases were randomly drawn with a probability of 0.5 from each of the two alleles of the genotype. Simulations that include reference bias first chose one allele as the reference. Bases were then drawn from this allele with probability *r* = 0.55.

Each base was substituted according to pre-defined error probabilities. These probabilities were derived from comparing the untreated Vindija 33.19 Neandertal data to the genotypes of the closely related Altai Neandertal. Separate substitution matrices were calculated for each of the first 15 and last 15 bases of sequences and one matrix for the remaining bases in the interior of sequences. Each simulated base was assigned one of the resulting 31 substitution matrices at random and was modified with the probabilities given by this matrix.

## 2.3 Ancient DNA data

I used several published datasets to test the performance of snpAD (Suppl. Table 1). Following previous approaches (Prüfer *et al.*, 2017), the analysis of all datasets was restricted to regions within a 35bp mapability track and sequences with MQ<25 and bases with Q<30 were removed. Some datasets consisted of libraries with different treatment that affect error rates. These types of libraries were considered separately for error estimation. Identical to the simulated data, 31 substitution matrices for the first 15 bases, the last 15 bases and central bases were estimated for each type of library.

## 2.3.1 Neandertal data for chromosome 21

I used published sequence data from chromosome 21 of the 30-fold coverage Vindija 33.19 genome (Prüfer *et al.*, 2017). Around 1/4 of this data was enzyme treated to remove ancient DNA damage, while the remaining data were not treated. For error estimation, treated and untreated data were regarded separately. In addition to the full dataset, the chromosome 21 data was subsampled to an average coverage of 3-25 using the samtools option “-s” (Li *et al.*, 2009).

Chromosome 21 has also been captured from sequencing libraries of a Neandertal sample from the El Sidrón Cave (Sid1253) and another Neandertal sample from the Vindija Cave (Vindija 33.15) (Kuhlwilm *et al.*, 2016). Note that Vindija 33.19 and 33.15 carry almost identical heterozygous sites on chromosome 21, suggesting that these two samples originate from the same Neandertal individual (Prüfer *et al.*, 2017).

## 2.3.2 Low-coverage Neandertals and modern humans

I used the recently published low-coverage genome sequences (1.0-fold to 2.7-fold coverage) for the late Neandertals Goyet, Spy, Vindija 87, Le Cotte and Mezmaiskaya 2 that are less than 50,000 years old (Hajdinjak *et al.*, 2018). For comparison with these low-coverage Neandertals, the genome-wide data of an untreated Vindija 33.19 library was subsampled to 0.9, 1.0, 1.2, 1.5 and 2.0-fold coverage and processed identically to other low-coverage samples.

In addition, I used 1-fold and 2-fold coverage subsamples from the 22-fold coverage genome of Loschbour, and the full data of Motala 12 (2.4-fold), both around 8000 year old modern human individuals from Europe (Lazaridis *et al.*, 2014).

## 3 Results

### 3.1 Assessing accuracy using simulated data

I first tested snpAD using simulated datasets of 10 million sites, each, ranging from 3 to 30-fold coverage. Parellelizing over 30 processor cores, individual simulations took between 23 and 83 minutes to process and under 6GiB of memory (see Suppl. Fig. 4 & 5).

Since the simulated genotype frequencies and the profile of simulated errors are known, the accuracy of inferred parameters can be estimated with respect to coverage. The simulations showed that a coverage of at least 4-fold is required to estimate genotype frequencies accurately (<10% deviation; Fig. 2). Parameter estimates for the simulation with 3-fold coverage deviated by 30-160% from simulated genotype probabilities (Suppl. Tables 2 & 3).

Lower coverage simulations contain a smaller number of observed bases. To exclude the possibility that a lack of informative sites explains the large deviations for the 3-fold coverage simulation, I repeated the analysis with 100 million sites at 3-fold coverage. As before, estimates deviated substantially from the simulated genotype probabilities (20-130%), indicating that lack of power is not the main reason for the deviation.

**Fig. 2.**
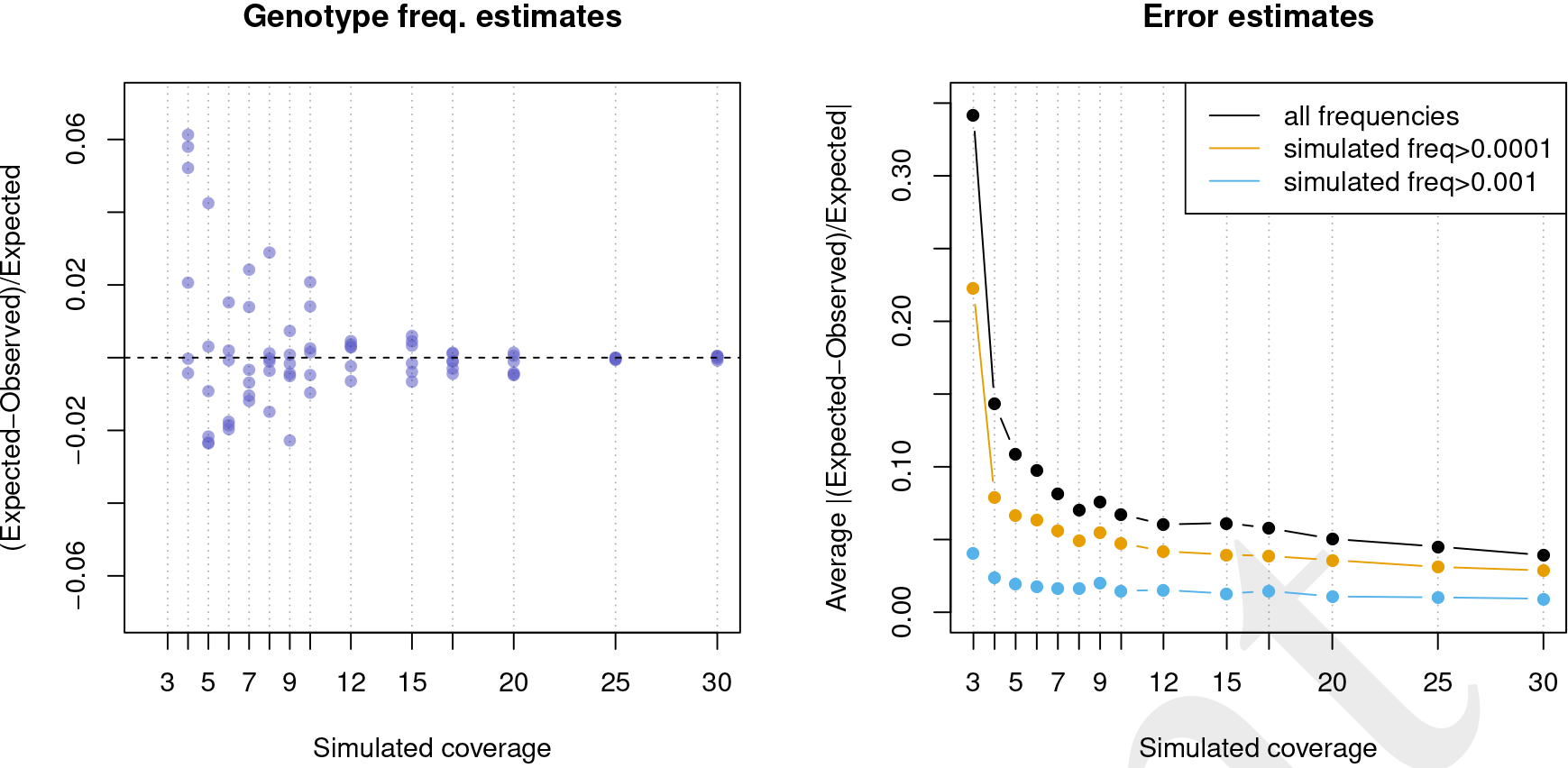
Accuracy of parameter estimation for simulated datasets. Left: Deviation from simulated genotype probabilities for the six heterozygous genotypes. Each simulation is indicated by a vertical blue dotted line and the estimates are shown as blue points. Estimates for 3-fold coverage deviated by more than 0.5 and are not visible at the depicted range. Right: Average deviation from simulated error probabilities.

Estimates of genotype frequencies were close to simulated parameters when the true error rates were given (<8% difference; Suppl. Table 5), indicating that error rates could not be estimated accurately at 3-fold coverage. Fig 2 shows that the estimated frequencies of errors are, on average, within 10% of the simulated frequencies for datasets with at least 6-fold coverage. Simulations with less than 6-fold coverage show larger deviation from the true parameters, especially for errors that occur at lower frequencies (Fig 2).

Note that these results are based on simulations with a fixed coverage. More realistic simulated coverages that follow a Poisson distribution (Lander and Waterman, 1988) yield more accurate genotype frequency estimates at lower coverage (Suppl. Table 6).

### 3.2 Subsamples of a high-coverage Neandertal genome

The Vindija 33.19 Neandertal was previously sequenced to 30-fold genomic coverage. The published genotypes were produced with an earlier version of snpAD, that required error rates to be specified. This error was estimated by comparing the Vindija 33.19 sequences to the closely related Altai Neandertal genome (Prüfer *et al.*, 2014). Genotype calls based on this approach were shown to outperform calls by GATK (Prüfer *et al.* 2017; see Suppl. Section 3 for a comparison of GATK with and without ancient DNA quality score rescaling (Jónsson *et al.*, 2013) to the latest snpAD version).

To test the accuracy of parameter estimates, I ran snpAD on the data for chromosome 21 of Vindija 33.19 (89 minutes wall clock runtime on 50 cores and ≈12GiB maximum memory usage). Since the true frequencies of error are not known, I used the differences of Vindija 33.19 sequences on chromosome 21 to the previously published Vindija 33.19 genotype calls as a baseline for comparison. The estimated error rates match this baseline well (Suppl. Fig. 6). The genotype frequencies show a difference of less than 1% from previous estimates (Suppl. Table 10).

Simulations indicated that snpAD performs well for high-coverage data, but that parameter estimates fit less well for coverage lower than 6-fold. To test whether these results also hold for a true ancient DNA dataset, I subsampled the chromosome 21 Vindija data to average coverages of 1 to 25-fold. SnpAD estimates on these subsampled data show that genotype and error rates are close (deviate by less than 10% on average) to the estimates with the full data (Fig. 3; Suppl. Table 11) as long as the average coverage is ≥15. Datasets with at least 2-fold coverage differed on average by at most 20% from the true genotype frequencies, and at most 30% from true error rates.

The estimated parameters can be used to determine the most likely genotypes along chromosome 21 for each subsampled dataset. To test how coverage affects the accuracy of the most likely call, these gentypes were compared to the genotypes gained from the full 30-fold coverage Vindija data (Suppl. Fig. 7 & 8; Suppl. Table 13). Less than 0.75% of calls at 1-fold coverage or higher were discordant, whereas datasets with at least 12.5-fold coverage showed less than 0.01%. Applying a cutoff on genotype quality scores (GQ30 or GQ50) further reduced the proportion of discordant calls (Suppl. Tables 14 & 15; Suppl. Figs. 9 & 10).

### 3.3 Reference bias

Capture and alignment procedures can introduce a bias in ancient sequence data that leads to an overrepresentation of sequences that support the capture bait or the reference genome used for alignment (reference bias). SnpAD supports the estimation of a parameter that captures this bias by testing for unequal representation of sequences supporting the reference and non-reference alleles at heterozygous sites.

Simulated data with a reference bias of 5% and 0% (*r* = 0.550 and *r* = 0.500) show that a minimum coverage of 15-fold is required to estimate reference bias (Suppl. Table 8). Simulations with this minimum coverage yield estimates of 0.549 − 0.558 for a simulated *r* = 0.550 and 0.500 − 0.508 for *r* = 0.500. That higher coverage is required to estimate reference bias is further corroborated by the subsampled Vindija 33.19 data, which fails to converge for simulations with ≤10-fold coverage and yields reference bias estimates from 7.8% to 4.7% for higher coverage datasets (Suppl. Table 12).

**Fig. 3.**
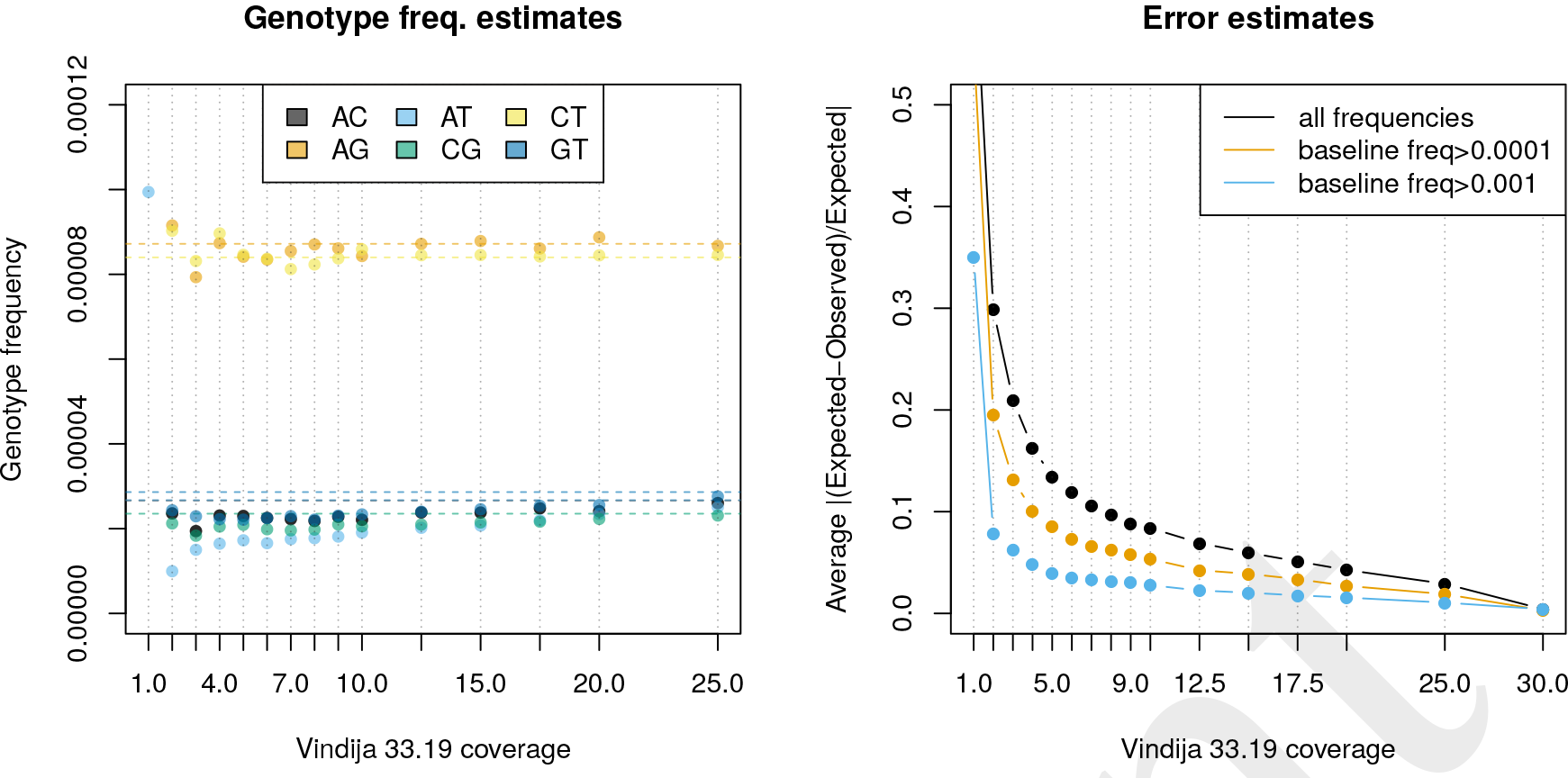
Parameter estimates for subsampled Vindija 33.19 data compared to full data. Left: Estimated genotype frequencies (points) compared to full data (shown as horizontal lines). Right: Average deviation from error rates in the full dataset. Estimates for 1-fold coverage fall outside of the plotted ranges.

Next, I ran snpAD for the chromosome 21 capture data of Vindija 33.15 and El Sidrón. Unfortunately, El Sidrón was too low coverage and parameter estimates did not converge. The higher-coverage Vindija 33.15 data, on the other hand, yielded an estimated 10% reference bias alongside similar genotype frequencies as Vindija 33.19 (Suppl. Table 16). A stronger reference bias for Vindija 33.15 than Vindija 33.19 could be explained by capture bias. This capture bias would favor sequences matching the capture bait, which, in the case of Vindija 33.15, was based on the human reference sequence.

To test whether genotype calling would benefit from incorporating reference bias, I called genotypes for Vindija 33.15 capture data with and without reference bias and compared these calls to those gained from Vindija 33.19 data (Suppl. Table 17). Note that Vindija 33.15 and 33.19 likely originate from the same individual so that genotype calls are expected to match. Vindija 33.15 calls with reference bias show a higher fraction of matching homozygous alternative genotypes (26157/41 match/do not match with bias, compared to 26117/81 without; Fisher’s exact test *p* = 0.0004) and matching heterozygous genotypes (3793/116 vs. 3785/124; *p* = 0.6464). However, the calls including reference bias also encompass more than 300 additional heterozygous calls that are not shared with Vindija 33.19. These results suggest that taking reference bias into account for genotype calling leads for Vindija 33.15 data to a larger fraction of false calls while gaining few additional sites that may be called correctly.

### 3.4 Low-coverage genomes

Results based on subsamples of the Vindija 33.19 data on chromosome 21 suggested that the power to estimate genotype frequencies and parameters of the error model are low for low-coverage samples. However, this analysis is limited by the small number of sites on chromosome 21. Furthermore, low-coverage shotgun data is expected to follow approximately a Poisson distribution (Lander and Waterman, 1988), so that even for low-coverage samples some fraction of sites exist that are covered much more often than the average. For a genome at an average 1-fold coverage, for instance, around 2% of sites are expected to be covered by at least 4 sequences (Suppl. Fig 11).

To test whether lower-coverage genomes can yield at least approximate estimates of heterozygosity, I subsampled a single library of Vindija 33.19 to between 0.9 and 2.0-fold coverage. This range of coverage is similar to the range observed in five recently published low-coverage Neandertal genomes ranging from 1.0 to 2.7-fold (Hajdinjak *et al.*, 2018). SnpAD was then run on all sites on the autosomes covered by at least four sequences and the estimates for genotype frequencies were compared to the genome-wide average for the high-coverage Vindija 33.19 Neandertal (Fig. 4). All low-coverage samples yielded overestimates of transition heterozygotes, although the difference to the high-coverage estimates grow smaller with increasing coverage. Trans version heterozygotes were in better agreement with expectation (maximum difference 17%; Suppl. Table 18).

To infer approximate estimates of heterozygosity, I ran snpAD on five low-coverage Neandertals and three low-coverage datasets of modern humans. Estimates of genotype frequencies indicate that all Neandertals are less heterozygous compared to the modern human data (Suppl. Table 19). Estimates for the Neandertals are generally close to estimates from Vindija 33.19, except for estimates for Spy, which are substantially higher (by 36-92%; Fig. 4). Among all tested Neandertals, the Spy individual is the sample with the lowest coverage and highest modern human contamination (1.7%), offering at least a partial explanation for the higher estimates.

### 3.5 Comparison with ATLAS

ATLAS, a software package for ancient DNA analyses, has been used to estimate heterozygosity from low-coverage genome data (Kousathanas *et al.*, 2017). Like other software (Lindgreen *et al.*, 2014; Zhou *et al.*, 2017), ATLAS uses a model of ancient DNA damage that expects rising rates of C to T exchanges towards the 5’-end and G to A exchanges at the 3’ end. Unfortunately, this model does not reflect the patterns of ancient DNA damage in sequencing libraries prepared with a more efficient single stranded protocol (Gansauge and Meyer, 2013; Meyer *et al.*, 2012) (Suppl. Figs 12 & 13). The latter protocol has been employed for the production of all Neandertal shotgun data used here for the evaluation of snpAD, so that snpAD and ATLAS cannot be compared on these datasets.

**Fig. 4.**
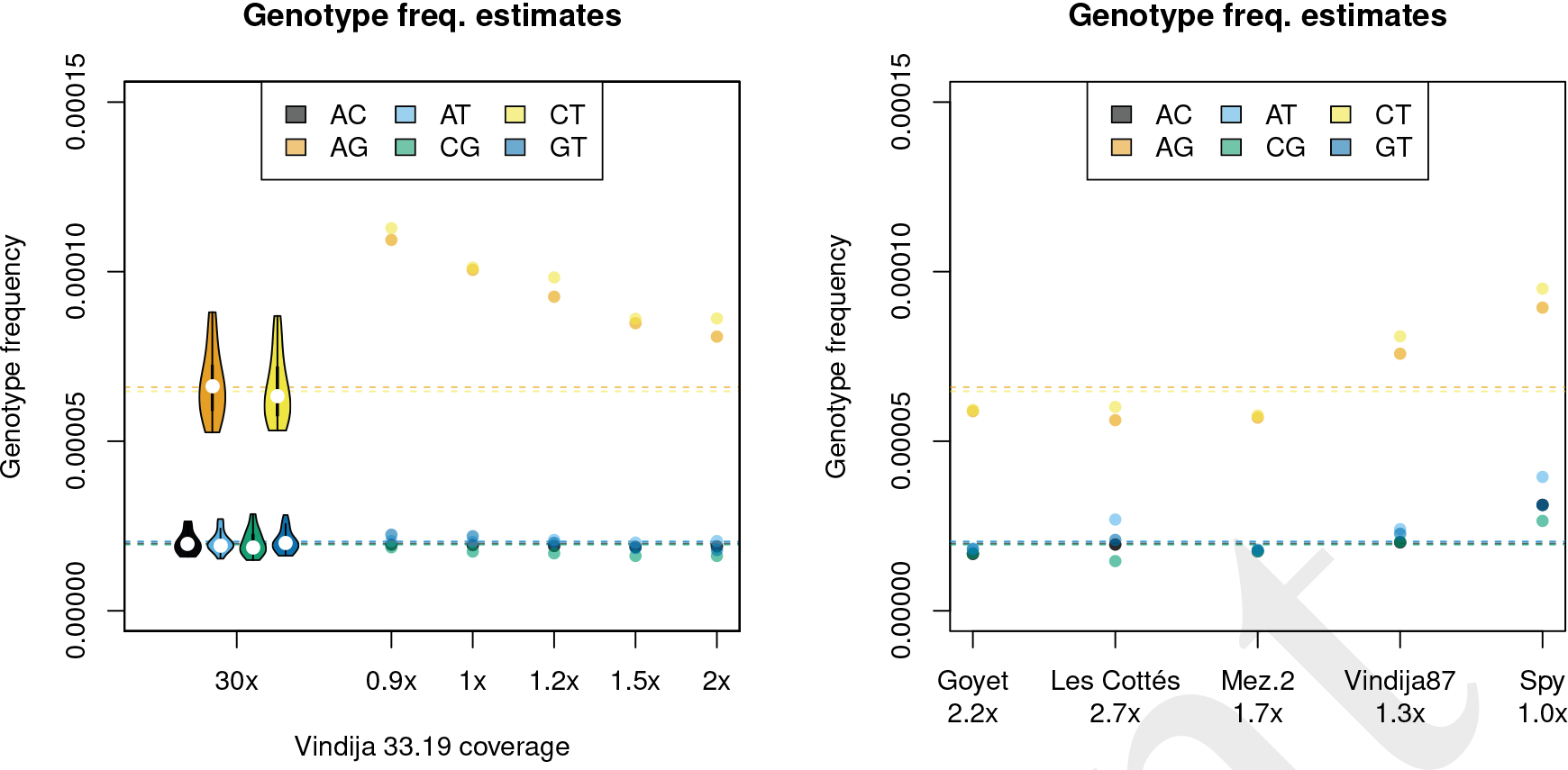
Genotype frequencies for autosomal sites with at least 4-fold coverage. Left: Vindija 33.19 full data and data from a single subsampled library. Violin plots show the distribution over Vindija 33.19 chromosomes. Right: Low-coverage Neandertals. Horizontal lines show genome-wide Vindija 33.19 estimate.

To make a comparison possible, I used the data of Motala12, an ≈8000 year old European hunter-gatherer (Lazaridis *et al.*, 2014) that was sequenced to to 2.4-fold coverage and matches ATLAS’ expected patterns of ancient DNA damage. ATLAS’ error estimates were based on conserved sites and heterozygosity estimates were determined using the *estimateTheta* function (see Suppl. Section 4 for details).

ATLAS estimated a heterozygosity of 1.24 × 10^−3^ whereas snpAD gave an estimate of 0.73 × 10^−3^ (Suppl. Tables 20 & 19). While the true heterozygosity of the individual is not known, the heterozygosity of the 22-fold coverage genome of Loschbour, also an ≈8000 year old hunter-gatherer from Europe, may serve as an proxy to put these numbers in perspective. Heterozygosity of Loschbour was estimated to be 0.66 × 10^−3^ using GATK (McKenna *et al.*, 2010; Lazaridis *et al.*, 2014) while an earlier version of snpAD yielded 0.62 × 10^−3^ (Prüfer *et al.*, 2017). I note that snpAD heterozygosity estimates of 1-fold and 2-fold subsamples (0.58 and 0.59 × 10^−3^, respectively) of the 22-fold Loschbour sample fall close to these estimates. Present-day non-Africans have been found to fall in the range of 0.5 − 0.7 × 10^−3^ (Mallick *et al.*, 2016).

## 4 Discussion/Conclusion

Calling genotypes from ancient DNA data is challenging due to the high errors rates in such data. Here, I showed that genotypes can be called by jointly estimating all necessary parameters from the data. Subsampling lower coverage subsets from a high coverage Neandertal genome indicated that the estimated parameters are reasonably accurate (<10% deviation) with at least 15-fold coverage. However, lower coverage data can still yield approximate estimates of heterozygosity.

The error model in snpAD differs from those implemented in other software for ancient DNA analyses. This model follows a minimalist approach, in that it only assumes that classes of bases exist that are characterized by the same error rates. The current implementation offers the option to classify observed bases by sequence position and type of library. However, the approach is extendable to incorporate other features that are informative of error rates. The simplistic design allows me to combine sequencing error and ancient DNA damage and to jointly estimate error rates and genotype frequencies. Furthermore, data with other types of error patterns than those observed in ancient DNA could be processed with snpAD without the need for adjustments to the model or subsequent estimation steps.

A perhaps underappreciated issue for ancient DNA analysis is reference bias (e.g. Prüfer *et al.*, 2010). Here I estimate this bias based on the overrepresentation of reference alleles at heterozygous positions. Using capture data from an Neandertal individual that has been shotgun sequenced to high-coverage, I was able to show that incorporating a uniform capture bias does not improve genotype calls. A shift of alleles at heterozygous sites also constitute part of the information used by estimators of modern human contamination in archaic individuals (Philip L.F. Johnson’s maximum likelihood estimator described in Prüfer *et al.*, 2014; Racimo *et al.*, 2016). The reference bias estimate thus provides an approximate upper limit for modern human contamination. Future work may aim to incorporate contamination estimates into the calling of genotypes to reconstruct sequences in the presence of contamination, similar in spirit to approaches to reconstruct mitochondrial genomes from contaminated sequence data (Renaud *et al.*, 2015).

To gain insight into the effective population sizes of late Neandertals, I used snpAD to estimate heterozygosity for five recently published low-coverage genomes (Hajdinjak *et al.*, 2018). While I caution that these estimates are approximate at best, it is intriguing that several late Neandertals yielded heterozygosity estimates that lie below those estimated from the high-coverage Vindija and Altai Neandertals. These results raise the possibility of particularly low heterozygosity in some of the late Neandertals, that could reflect a small number of individuals towards the end of the Neandertal’s reign in Europe.

## Acknowledgements

I would like to thank Nick Patterson for valuable input that motivated further improvements, and the members of the genomics, bioinformatics and ancient DNA groups at the Max Planck Institute for interesting
discussions. I am indepted to Cesare de Filippo who helped with earlier tests of ATLAS and snpAD, Michael Dannemann and Fabrizio Mafessoni for critically reading the manuscript, and all members of the Vindija Genome Analysis consortium for patience with and discussions of snpAD genotype calls.

## Funding

This work has been supported by the Max Planck Society, the Max-Planck-Förderstiftung (grant 31-12LMP Pääbo), the Strategischer Innovationsfonds der Max-Planck-Gesellschaft and the European Research Council (ERC) (grant agreement no. 694707).

