## Supplementary Materials for "snpAD: An ancient DNA genotype caller" (from another version of this preprint: version 1, 2018-03-24)

### snpAD: an ancient DNA genotyper

#### Supplementary Tables and Figures

March 24, 2018

#### Contents

1 Figures 4

2 Tables 14

3 Comparison to ATLAS 28

10 Estimated genotype frequencies for subsampled Vindija 33.19 data 19
11 Estimated genotype frequencies for subsampled Vindija 33.19 data with reference bias . . . . 20
12 Comparison of called genotypes in Vindija 33.19 subsamples to 30x calls. . . . . 21
13 Comparison of called genotypes at GQ30 in Vindija 33.19 subsamples to 30x calls. . . . . 22

14 Comparison of called genotypes at GQ50 in Vindija 33.19 subsamples to 30x calls. . . . 23
15 Vindija 33.15: genotype frequency estimates with and without reference bias . . . 24
16 Comparison of Vindija 33.19 and 33.15 chr21 genotypes with and without reference bias . . . 25
17 Genotype frequencies for subsampled Vindija 33.19 data restricting to at least 4-fold coverage . . . 26
18 Genotype frequencies for low-coverage archaic and modern humans with at least 4-fold coverage . . . 27
19 ATLAS' genome-wide estimates of $\theta$ for Motala12 . . . 28

### 1 Figures

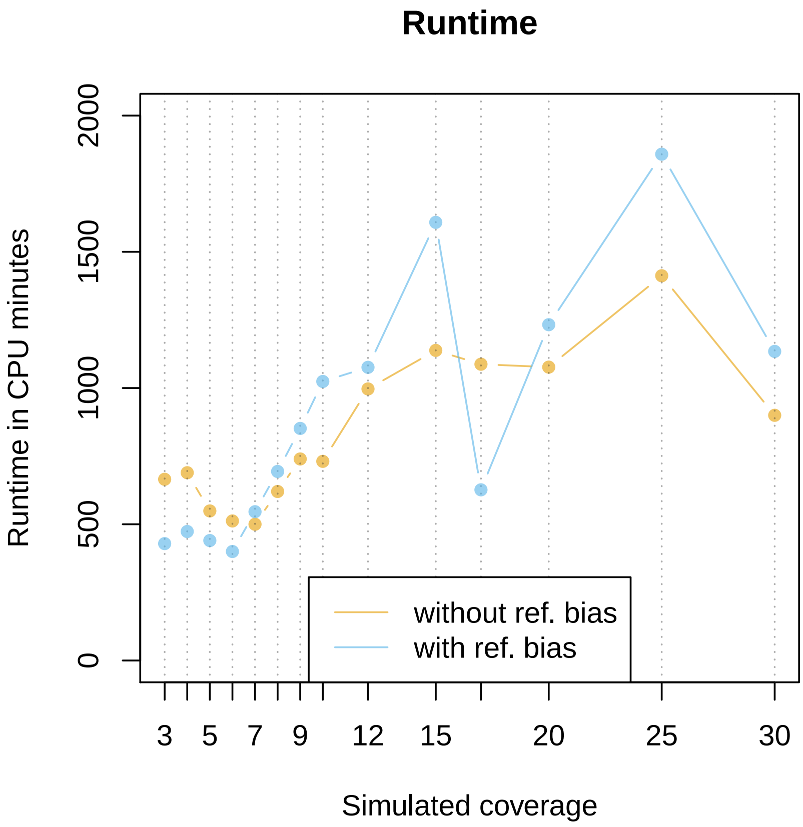

Figure 1: Runtime in CPU minutes for estimates with and without reference bias on simulated datasets. Wall clock time ranged from 23 to 83 minutes parellelizing over 30 cores of a multi-core Intel Xeon server.

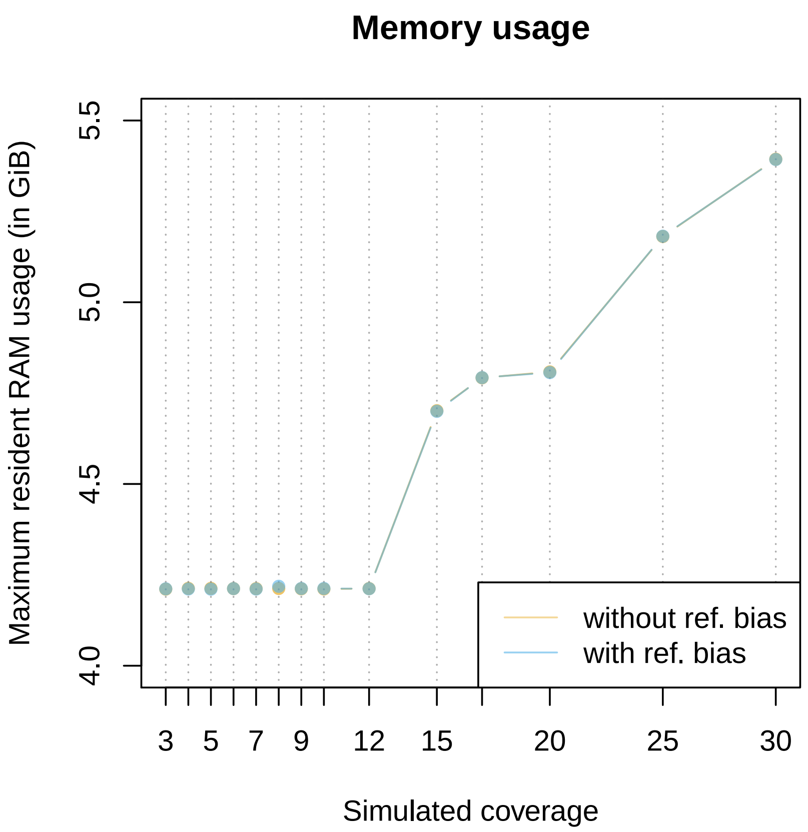

Figure 2: Maximum RAM usage in GiB during estimation of parameters with and without reference bias on simulated datasets. Note that the estimates overlap almost perfectly between the runs with and without reference bias so that points appear in light gray. Y-axis starts at 4GiB.

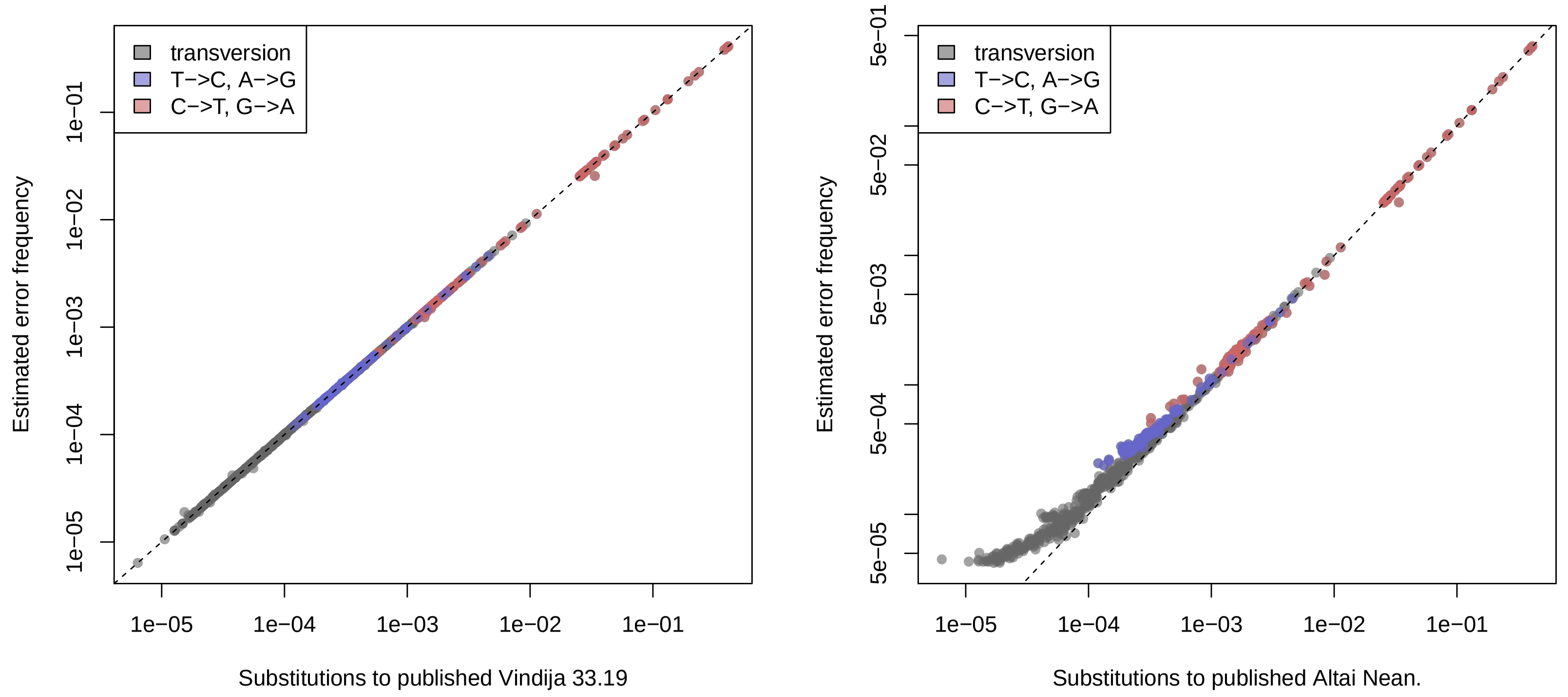

Figure 3: Error estimates for Vindija 33.19 data on chromosome 21. Left: Error estimates compared to substitutions of the Vindija 33.19 sequences to the previously published Vindija 33.19 genotypes. Right: Error estimates compared to substitutions of the data to the Altai Neandertal genotypes.

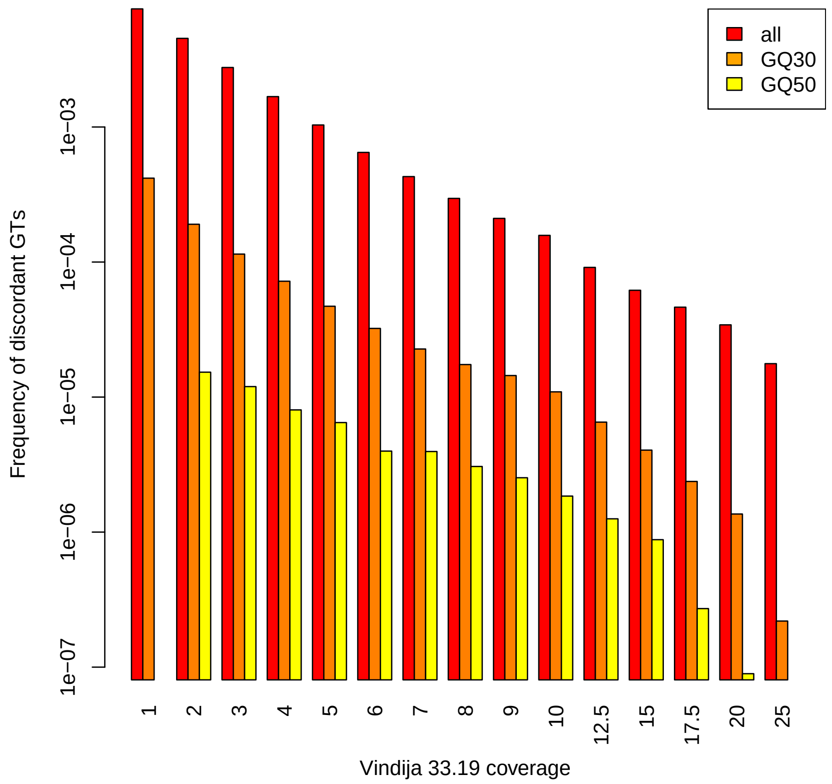

Figure 4: Discordant genotype calls in Vindija 33.19 subsamples compared to the full 30x coverage calls. Frequency of discordant genotypes is on a log-scale. Colors indicate whether all genotypes were considered, or only those with a genotype quality >30 or >50.

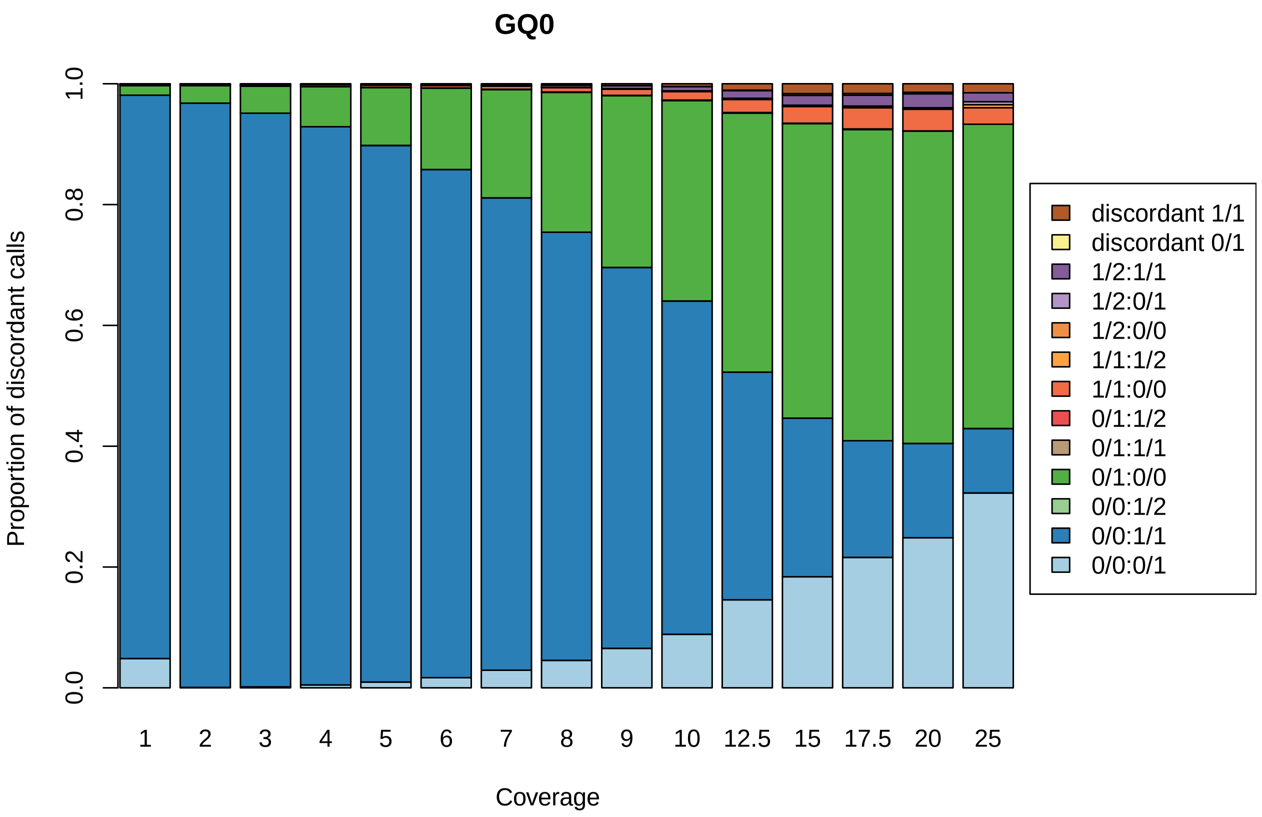

Figure 5: Classification of discordant genotype calls in Vindija 33.19 subsamples. No genotype quality cutoff was applied. For counts see Table 12

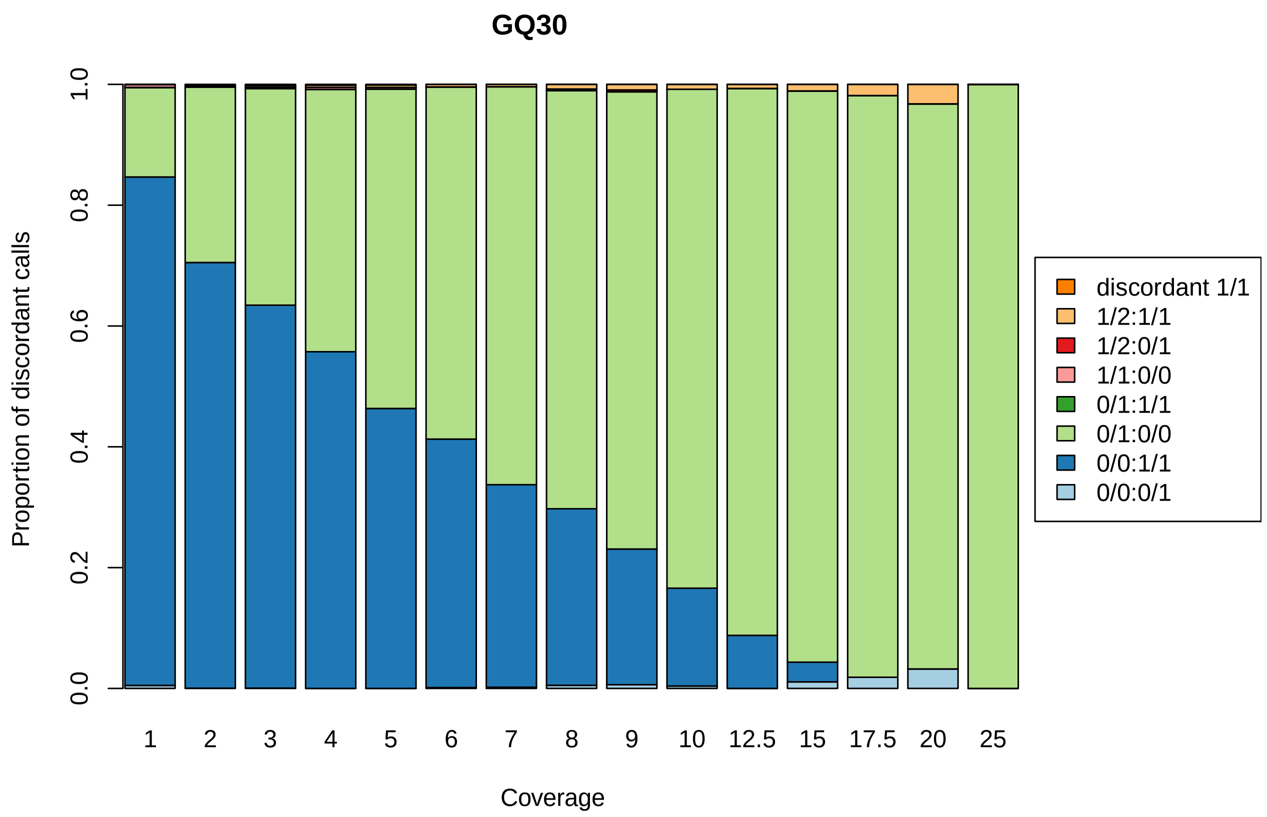

Figure 6: Classification of discordant genotype calls in Vindija 33.19 subsamples. A genotype quality cutoff of $\geq 30$ was applied. For counts see Table 12

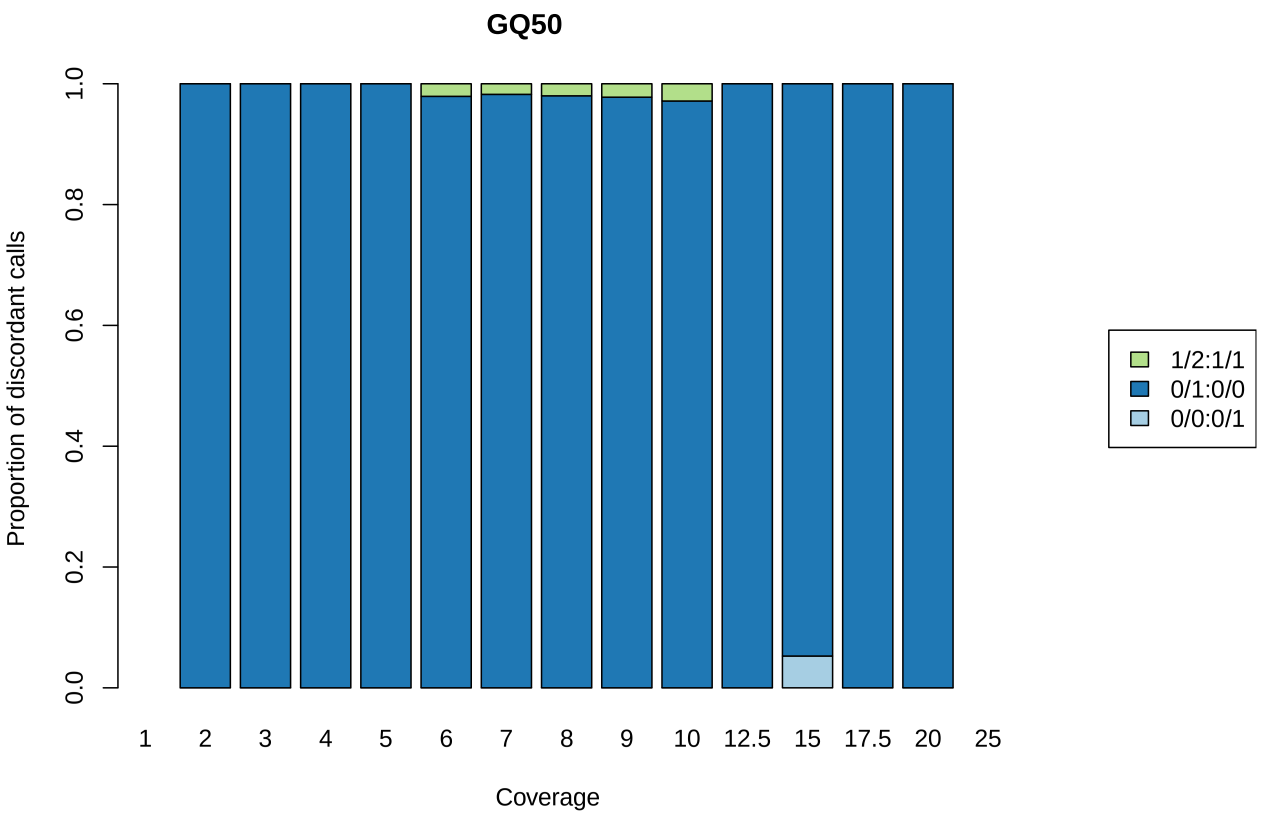

Figure 7: Classification of discordant genotype calls in Vindija 33.19 subsamples. A genotype quality cutoff of ≥50 was applied. No discordant calls were observed for coverage 1 and 25. For counts see Table 12

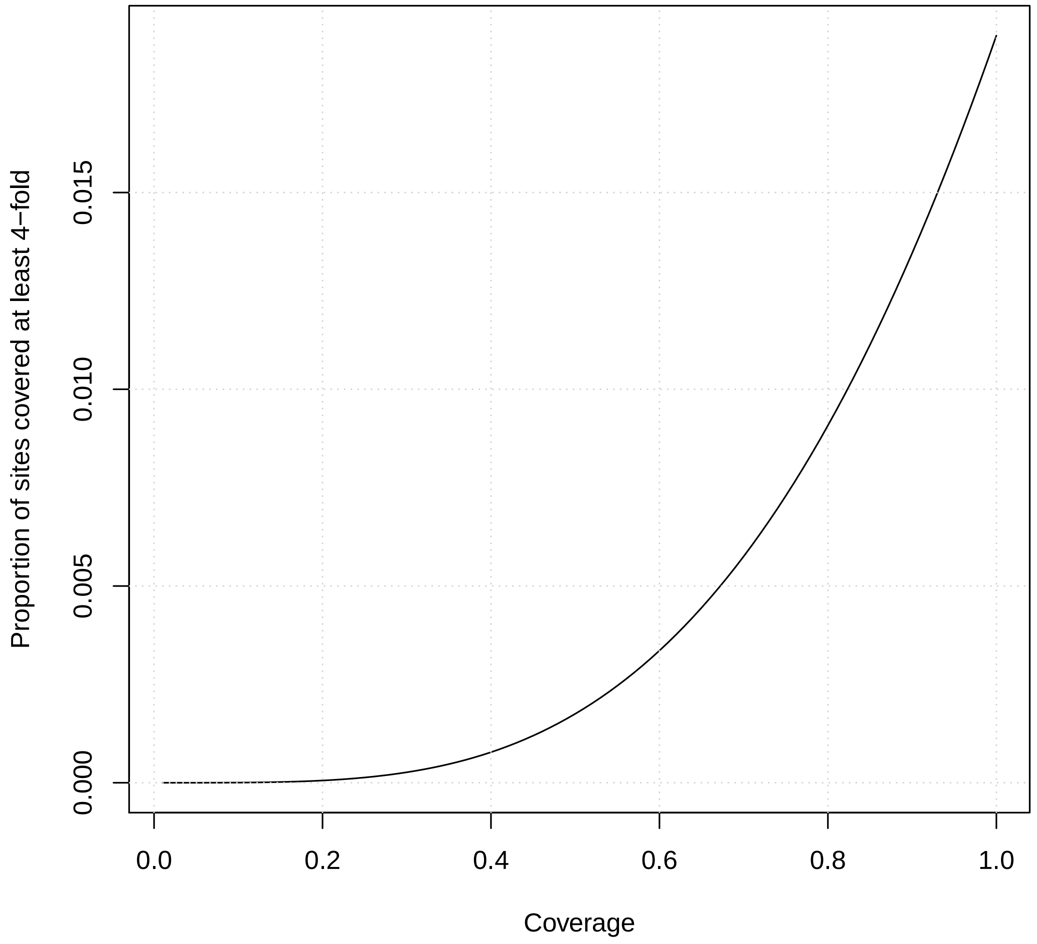

Figure 8: Expected proportion of sites with at least 4-fold coverage in low-coverage genomes according to the Lander-Waterman statistics.

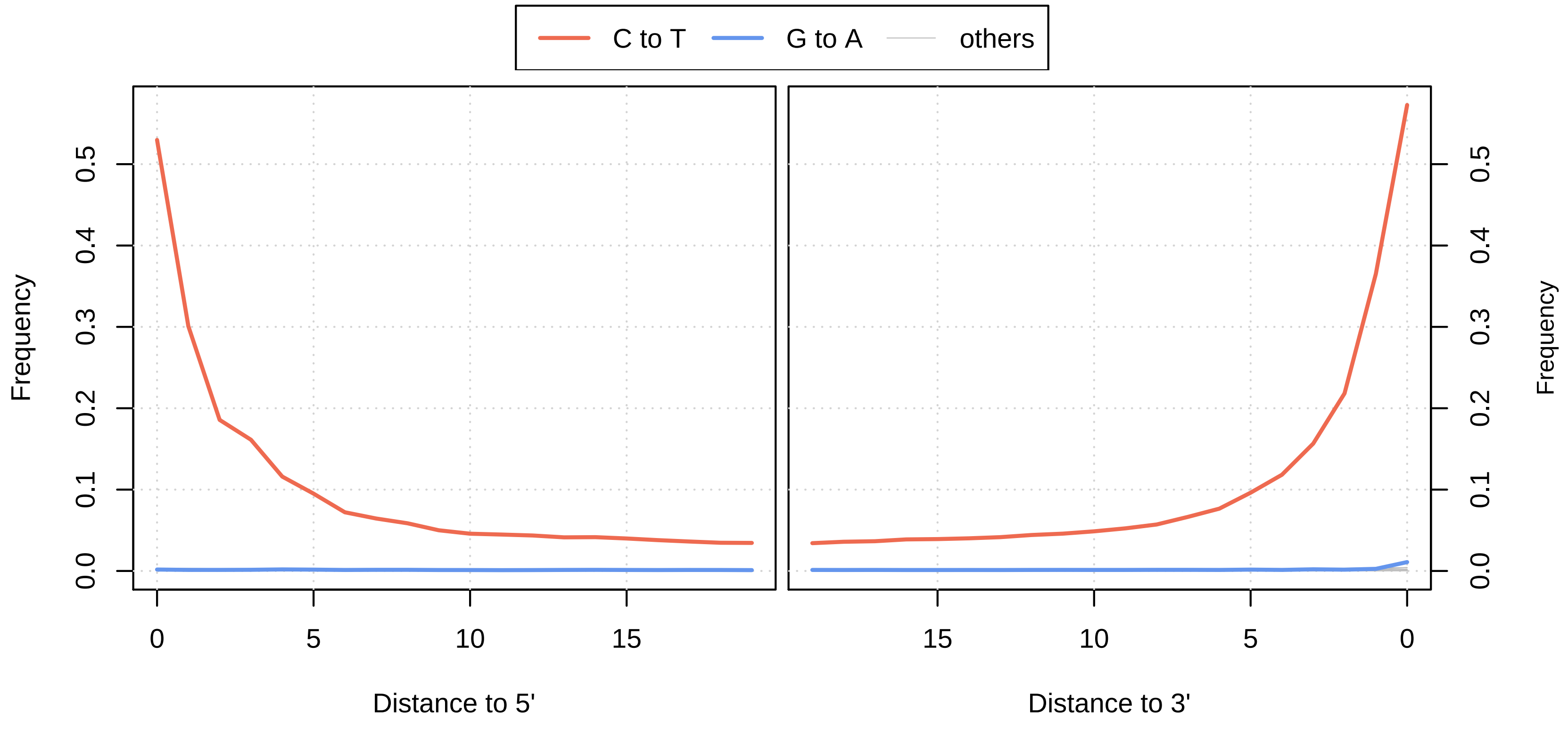

Figure 9: Damage patterns in the Vindija 87 Neandertal data prepared with a single stranded library protocol.

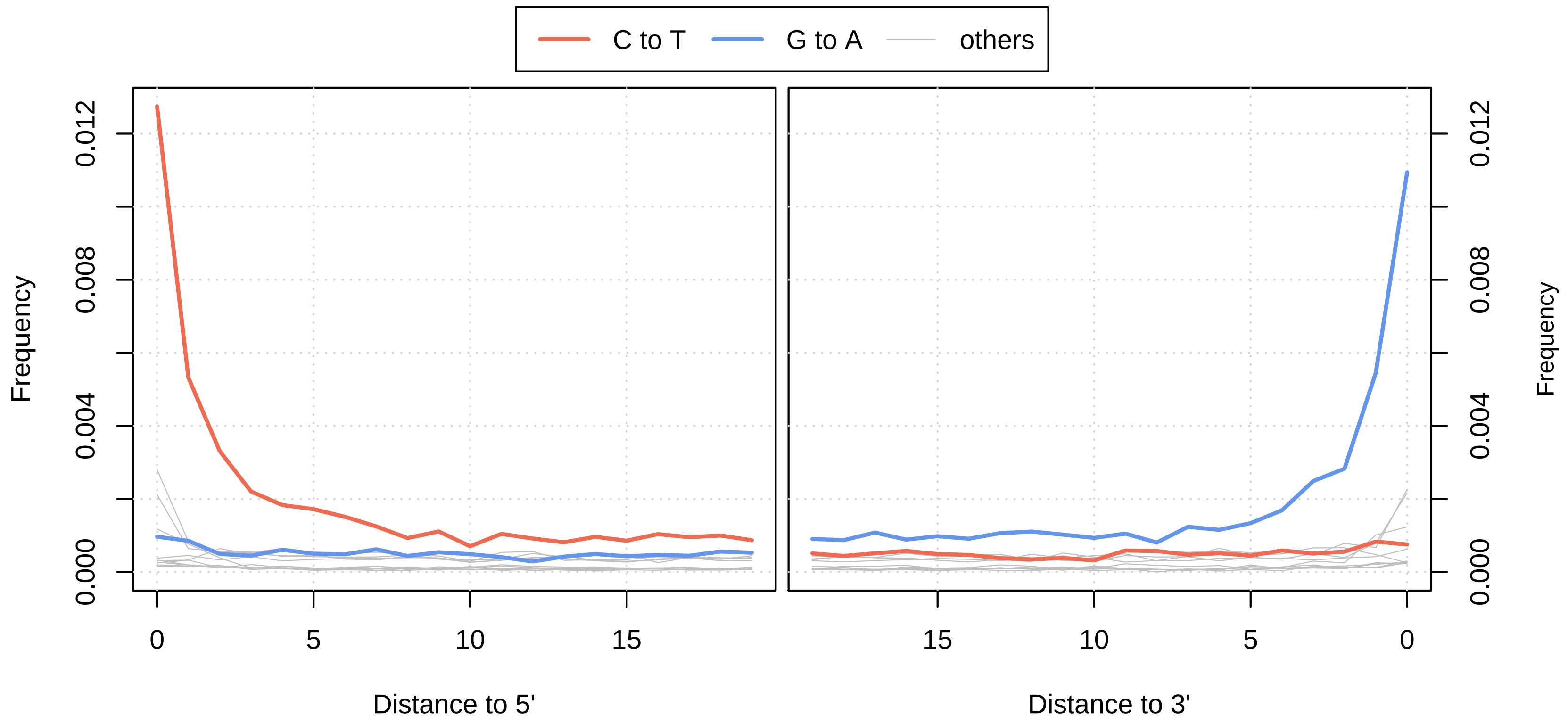

Figure 10: Damage patterns in the data of the European hunter-gatherer Motala12 prepared with a double stranded library protocol.

### 2 Tables

| Group | Name | Coverage | Reference |
|---|---|---|---|
| Neandertal | Altai (from Denisova cave) | 52x | Prüfer *et al.* (2014) |
| Neandertal | Vindija 33.19 | 30x | Prüfer *et al.* (2017) |
| Modern human | Loschbour | 22x | Lazaridis *et al.* (2014) |
| Neandertal | Les Cottés Z4-1514 | 2.7x | Hajdinjak *et al.* (2018) |
| Modern human | Motala 12 | 2.4x | Lazaridis *et al.* (2014) |
| Neandertal | Goyet Q56-1 | 2.2x | Hajdinjak *et al.* (2018) |
| Neandertal | Mezmaiskaya 2 | 1.7x | Hajdinjak *et al.* (2018) |
| Neandertal | Vindija 87 | 1.3x | Hajdinjak *et al.* (2018) |
| Neandertal | Spy 94a | 1.0x | Hajdinjak *et al.* (2018) |

Table 1: Datasets used in this study.

| Cov. | AC | AG | AT | CG | CT | GT |
|---|---|---|---|---|---|---|
| 3 | 0.0994 | 0.193 | 0.0979 | 0.0958 | 0.197 | 0.0977 |
| 4 | 0.0983 | 0.2045 | 0.1008 | 0.0977 | 0.2074 | 0.1056 |
| 5 | 0.0987 | 0.2035 | 0.105 | 0.1006 | 0.2051 | 0.092 |
| 6 | 0.1001 | 0.1973 | 0.1002 | 0.103 | 0.2035 | 0.0975 |
| 7 | 0.0998 | 0.2004 | 0.1001 | 0.1024 | 0.2083 | 0.1037 |
| 8 | 0.103 | 0.1964 | 0.1007 | 0.1014 | 0.1985 | 0.0961 |
| 9 | 0.0966 | 0.2057 | 0.0916 | 0.0972 | 0.1986 | 0.095 |
| 10 | 0.1039 | 0.1952 | 0.096 | 0.1 | 0.2053 | 0.1003 |
| 12 | 0.1051 | 0.2022 | 0.1 | 0.0949 | 0.2033 | 0.0989 |
| 15 | 0.0994 | 0.1985 | 0.1009 | 0.099 | 0.2003 | 0.1027 |
| 17 | 0.1014 | 0.2019 | 0.0992 | 0.0993 | 0.2026 | 0.0964 |
| 20 | 0.099 | 0.1939 | 0.1018 | 0.0998 | 0.2012 | 0.1031 |
| 25 | 0.1045 | 0.2009 | 0.0971 | 0.0964 | 0.2007 | 0.0986 |
| 30 | 0.0976 | 0.2051 | 0.1019 | 0.0974 | 0.2011 | 0.0976 |

Table 2: Simulated genotype frequencies (per 1000bp). Deviation from specified parameters are due to the use of a pseudo random number generator during simulation.

| Cov. | AC | AG | AT | CG | CT | GT |
|---|---|---|---|---|---|---|
| 3 | 0.141943 | 0.487617 | 0.132684 | 0.217547 | 0.518872 | 0.145262 |
| 4 | 0.0978802 | 0.217037 | 0.106058 | 0.103366 | 0.207338 | 0.107778 |
| 5 | 0.0977911 | 0.19909 | 0.102525 | 0.0982538 | 0.213814 | 0.0922814 |
| 6 | 0.0982396 | 0.2003 | 0.100129 | 0.100975 | 0.2039 | 0.0957835 |
| 7 | 0.0994662 | 0.205249 | 0.099413 | 0.101335 | 0.211202 | 0.102465 |
| 8 | 0.102891 | 0.202079 | 0.100336 | 0.101366 | 0.195546 | 0.0962143 |
| 9 | 0.0961095 | 0.201015 | 0.0916744 | 0.0967756 | 0.198276 | 0.0956978 |
| 10 | 0.102898 | 0.195699 | 0.0955418 | 0.10141 | 0.209569 | 0.100444 |
| 12 | 0.105482 | 0.200895 | 0.100301 | 0.0951686 | 0.204228 | 0.0986684 |
| 15 | 0.0998465 | 0.199688 | 0.100235 | 0.0986095 | 0.199992 | 0.10304 |
| 17 | 0.101299 | 0.202168 | 0.0993153 | 0.0990021 | 0.201701 | 0.0962989 |
| 20 | 0.0985454 | 0.194164 | 0.101311 | 0.099694 | 0.200358 | 0.103154 |
| 25 | 0.104501 | 0.200799 | 0.0970826 | 0.0963994 | 0.200622 | 0.0985396 |
| 30 | 0.0976482 | 0.20494 | 0.101913 | 0.0973725 | 0.201186 | 0.0976336 |

Table 3: Estimated genotype frequencies (per 1000bp) from simulations (see Table 2).

| Genotype | Simulated freq. | Estimated freq. | Exp-Obs/Exp |
|---|---|---|---|
| AC | 0.10027 | 0.132199 | -0.318 |
| AG | 0.19781 | 0.242159 | -0.224 |
| AT | 0.10025 | 0.020440 | 0.7961 |
| CG | 0.10238 | 0.230882 | -1.255 |
| CT | 0.19831 | 0.235709 | -0.189 |
| GT | 0.10012 | 0.131489 | -0.313 |

Table 4: Estimated genotype frequencies (per 1000bp) from a simulation with 100 million sites at exactly 3-fold coverage.

| Genotype | Simulated freq. | Estimated freq. with true error rates | Exp-Obs/Exp |
|---|---|---|---|
| AC | 9.94e-05 | 9.67491e-05 | 0.026669014084507 |
| AG | 0.000193 | 0.000178968 | 0.0727046632124353 |
| AT | 9.79e-05 | 9.88342e-05 | -0.00954239019407561 |
| CG | 9.58e-05 | 9.81046e-05 | -0.0240563674321503 |
| CT | 0.000197 | 0.000211775 | -0.0750000000000001 |
| GT | 9.77e-05 | 0.000102151 | -0.0455578300921188 |

Table 5: Genotype frequency estimates for 3x simulated coverage when true error rates are given.

| Cov. | AC | AG | AT | CG | CT | GT | ref.bias |
|---|---|---|---|---|---|---|---|
| 3 | 0.1071 | 0.2018 | 0.0989 | 0.094 | 0.1988 | 0.104 | 0.5511 |
| 4 | 0.0983 | 0.1951 | 0.1047 | 0.1017 | 0.199 | 0.098 | 0.5494 |
| 5 | 0.0983 | 0.1975 | 0.0975 | 0.0969 | 0.1965 | 0.1011 | 0.5461 |
| 6 | 0.096 | 0.203 | 0.0977 | 0.0977 | 0.1991 | 0.0996 | 0.5498 |
| 7 | 0.0993 | 0.1914 | 0.0976 | 0.1003 | 0.1969 | 0.0972 | 0.5464 |
| 8 | 0.0997 | 0.1981 | 0.0993 | 0.0992 | 0.201 | 0.0998 | 0.5475 |
| 9 | 0.1027 | 0.1996 | 0.0977 | 0.105 | 0.2007 | 0.097 | 0.5510 |
| 10 | 0.0958 | 0.2 | 0.0991 | 0.1047 | 0.2023 | 0.0966 | 0.5515 |
| 12 | 0.0976 | 0.2029 | 0.0985 | 0.0987 | 0.1966 | 0.0995 | 0.5508 |
| 15 | 0.0962 | 0.1955 | 0.0966 | 0.1 | 0.2036 | 0.1045 | 0.5501 |
| 17 | 0.1018 | 0.1918 | 0.0966 | 0.0953 | 0.202 | 0.0966 | 0.5503 |
| 20 | 0.1019 | 0.1963 | 0.0992 | 0.1039 | 0.1928 | 0.0958 | 0.5485 |
| 25 | 0.0964 | 0.2025 | 0.0946 | 0.0986 | 0.2072 | 0.1016 | 0.5488 |
| 30 | 0.0996 | 0.2049 | 0.1047 | 0.1009 | 0.1997 | 0.1044 | 0.5514 |

Table 6: Simulated genotype frequencies with reference bias (per 1000bp).

| Cov. | AC | AG | AT | CG | CT | GT | ref.bias |
|---|---|---|---|---|---|---|---|
| 3 | 1 | 1 | 1 | 1 | 1 | 1 | 0.95417 |
| 4 | 1 | 1 | 1 | 1 | 1 | 1 | 0.953826 |
| 5 | 1 | 1 | 1 | 1 | 1 | 1 | 0.956578 |
| 6 | 1 | 1 | 1 | 1 | 1 | 1 | 0.958906 |
| 7 | 1 | 1 | 1 | 1 | 1 | 1 | 0.958269 |
| 8 | 1 | 1 | 1 | 1 | 1 | 1 | 0.953793 |
| 9 | 1 | 1 | 1 | 1 | 1 | 1 | 0.950293 |
| 10 | 0.103956 | 0.251935 | 0.107054 | 0.115078 | 0.252771 | 0.104944 | 0.608715 |
| 12 | 0.0993383 | 0.22121 | 0.0997244 | 0.101436 | 0.21645 | 0.102138 | 0.5730837 |
| 15 | 0.0977169 | 0.202107 | 0.0976777 | 0.100955 | 0.210079 | 0.104663 | 0.558126 |
| 17 | 0.101805 | 0.197107 | 0.0978276 | 0.096013 | 0.203464 | 0.0969439 | 0.5546397 |
| 20 | 0.101403 | 0.198897 | 0.0989341 | 0.104611 | 0.195179 | 0.0956663 | 0.5505485 |
| 25 | 0.0970535 | 0.201382 | 0.0935319 | 0.0976879 | 0.20715 | 0.101026 | 0.549275 |
| 30 | 0.0996463 | 0.20537 | 0.104621 | 0.101775 | 0.202138 | 0.104083 | 0.5518523 |

Table 7: Estimated genotype frequencies (per 1000bp) and reference bias from simulations with reference bias (see Table 6). Frequencies of 1/1000bp correspond to the initial values for the optimization procedure and indicate that the algorithm did not converge.

| Cov. | AC | AG | AT | CG | CT | GT | ref.bias |
|---|---|---|---|---|---|---|---|
| 3 | 1 | 1 | 1 | 1 | 1 | 1 | 0.953438 |
| 4 | 1 | 1 | 1 | 1 | 1 | 1 | 0.953484 |
| 5 | 1 | 1 | 1 | 1 | 1 | 1 | 0.956395 |
| 6 | 1 | 1 | 1 | 1 | 1 | 1 | 0.958075 |
| 7 | 1 | 1 | 1 | 1 | 1 | 1 | 0.956719 |
| 8 | 1 | 1 | 1 | 1 | 1 | 1 | 0.95392 |
| 9 | 0.103322 | 0.237223 | 0.0982436 | 0.104155 | 0.234996 | 0.103418 | 0.5754379 |
| 10 | 0.105007 | 0.207003 | 0.0977101 | 0.103477 | 0.22399 | 0.103373 | 0.5405103 |
| 12 | 0.10596 | 0.208124 | 0.0997445 | 0.0944469 | 0.20766 | 0.100264 | 0.518081 |
| 15 | 0.100901 | 0.20544 | 0.100627 | 0.0985946 | 0.197782 | 0.102163 | 0.50840239 |
| 17 | 0.10114 | 0.202382 | 0.0990932 | 0.0991423 | 0.202261 | 0.0962275 | 0.50315541 |
| 20 | 0.0986462 | 0.194231 | 0.102811 | 0.0999103 | 0.200908 | 0.103067 | 0.5000655172 |
| 25 | 0.104676 | 0.19986 | 0.0975324 | 0.0960198 | 0.201889 | 0.0984695 | 0.50172859 |
| 30 | 0.0975958 | 0.204819 | 0.10192 | 0.0974037 | 0.201259 | 0.0976113 | 0.50130228 |

Table 8: Estimated genotype frequencies (per 1000bp) and reference bias from simulations without reference bias (see Table 2). Simulations included no reference bias ($r = 0.50$), but fluctuated between 0.498-0.502 due to the use of a pseudo random number generator.

| Genotype | Previous estimates | Current estimates | Exp-Obs/Exp |
|---|---|---|---|
| AC | 0.02637483448 | 0.0266535 | -0.0106 |
| AG | 0.08703966695 | 0.0871984 | -0.0018 |
| AT | 0.02642543445 | 0.0267161 | -0.0110 |
| CG | 0.02333611154 | 0.0235279 | -0.0082 |
| CT | 0.08398376852 | 0.0839987 | -0.0002 |
| GT | 0.02822805388 | 0.028621 | -0.0139 |

Table 9: Vindija 33.19: previous and new genotype frequency estimates for 30x chromosome 21. Previous estimates were based on an error rates derived from comparing Vindija 33.19 sequences to the Altai Neandertal genotypes, while the new procedure co-estimates the error. Genotype frequencies are given per 1000bp.

| Cov. | AC | AG | AT | CG | CT | GT |
|---|---|---|---|---|---|---|
| 1 | 0.156481 | 1 | 0.099432 | 0.181886 | 1 | 0.148702 |
| 2 | 0.0235957 | 0.0915316 | 0.00995462 | 0.0212318 | 0.0902833 | 0.0243273 |
| 3 | 0.0194142 | 0.0793266 | 0.015004 | 0.0183244 | 0.0831413 | 0.0229132 |
| 4 | 0.0230869 | 0.0873542 | 0.0164465 | 0.0204864 | 0.0896604 | 0.0222243 |
| 5 | 0.022984 | 0.0841431 | 0.017246 | 0.0208929 | 0.0846063 | 0.0221146 |
| 6 | 0.0225169 | 0.0834912 | 0.016566 | 0.0198478 | 0.0836662 | 0.022483 |
| 7 | 0.0222535 | 0.0854938 | 0.0175251 | 0.0195863 | 0.0812381 | 0.022945 |
| 8 | 0.0218176 | 0.087055 | 0.0177506 | 0.0197764 | 0.0823211 | 0.0221323 |
| 9 | 0.0228057 | 0.0861142 | 0.0181263 | 0.0209766 | 0.0837547 | 0.0230667 |
| 10 | 0.0220717 | 0.0843099 | 0.0190784 | 0.0205518 | 0.0859023 | 0.0233493 |
| 12.5 | 0.0238188 | 0.0871949 | 0.0201923 | 0.0209605 | 0.0845151 | 0.0239949 |
| 15 | 0.0237837 | 0.0878583 | 0.020731 | 0.0214662 | 0.0845611 | 0.0245654 |
| 17.5 | 0.0247929 | 0.0860958 | 0.0219419 | 0.0216504 | 0.0842108 | 0.0253737 |
| 20 | 0.0241059 | 0.0887325 | 0.0234373 | 0.0222888 | 0.0844751 | 0.0255594 |
| 25 | 0.0260217 | 0.0867229 | 0.0250622 | 0.0230925 | 0.0845706 | 0.0275897 |
| 30 | 0.0266535 | 0.0871984 | 0.0267161 | 0.0235279 | 0.0839987 | 0.028621 |

Table 10: Estimated genotype frequencies (per 1000bp) for subsampled Vindija 33.19 data and full data (30-fold coverage) for Chromosome 21.

| Cov. | AC | AG | AT | CG | CT | GT | ref.bias. |
|---|---|---|---|---|---|---|---|
| 1 | 1 | 1 | 1 | 1 | 1 | 1 | 0.488789 |
| 2 | 1 | 1 | 1 | 1 | 1 | 1 | 0.483124 |
| 3 | 1 | 1 | 1 | 1 | 1 | 1 | 0.482451 |
| 4 | 1 | 1 | 1 | 1 | 1 | 1 | 0.483107 |
| 5 | 1 | 1 | 1 | 1 | 1 | 1 | 0.483965 |
| 6 | 1 | 1 | 1 | 1 | 1 | 1 | 0.484572 |
| 7 | 1 | 1 | 1 | 1 | 1 | 1 | 0.484614 |
| 8 | 1 | 1 | 1 | 1 | 1 | 1 | 0.484388 |
| 9 | 1 | 1 | 1 | 1 | 1 | 1 | 0.48411 |
| 10 | 1 | 1 | 1 | 1 | 1 | 1 | 0.48357 |
| 12.5 | 0.0254813 | 0.102948 | 0.0211019 | 0.0233485 | 0.0974866 | 0.0260767 | 0.0780759 |
| 15 | 0.0250461 | 0.095778 | 0.0220548 | 0.0220069 | 0.0927085 | 0.0257213 | 0.0641697 |
| 17.5 | 0.0255528 | 0.0917879 | 0.0226941 | 0.0221606 | 0.0896427 | 0.0262666 | 0.055925 |
| 20 | 0.026011 | 0.0911584 | 0.0243382 | 0.0229291 | 0.0882428 | 0.0274094 | 0.0526592 |
| 25 | 0.0265753 | 0.0896112 | 0.0260777 | 0.0234638 | 0.0872355 | 0.0283603 | 0.0495509 |
| 30 | 0.0271757 | 0.0892411 | 0.0278416 | 0.0238781 | 0.086252 | 0.0291206 | 0.0471163 |

Table 11: Estimated genotype frequencies (per 1000bp) and reference bias for subsampled Vindija 33.19 data and full data (30-fold coverage) for Chromosome 21.

| Genotypes | 1x | 2x | 3x | 4x | 5x | 6x | 7x | 8x | 9x | 10x | 12.5x | 15x | 17.5x | 20x | 25x |
|---|---|---|---|---|---|---|---|---|---|---|---|---|---|---|---|
| 0/0:0/1 | 5152 | 63 | 94 | 180 | 222 | 247 | 284 | 306 | 313 | 318 | 303 | 259 | 228 | 194 | 130 |
| 0/0:1/1 | 99102 | 84857 | 55902 | 34348 | 20669 | 12343 | 7623 | 4773 | 3022 | 1981 | 784 | 370 | 204 | 122 | 43 |
| 0/0:1/2 | 8 | 0 | 0 | 0 | 1 | 0 | 0 | 0 | 0 | 0 | 0 | 0 | 0 | 0 | 0 |
| 0/1:0/0 | 1711 | 2542 | 2632 | 2469 | 2230 | 1975 | 1748 | 1559 | 1362 | 1191 | 892 | 687 | 544 | 404 | 203 |
| 0/1:1/1 | 19 | 15 | 13 | 8 | 8 | 5 | 4 | 4 | 3 | 3 | 2 | 1 | 1 | 0 | 0 |
| 0/1:1/2 | 2 | 0 | 0 | 0 | 0 | 0 | 0 | 0 | 0 | 0 | 0 | 0 | 0 | 0 | 0 |
| 1/1:0/0 | 197 | 192 | 149 | 108 | 89 | 64 | 51 | 49 | 50 | 51 | 45 | 39 | 37 | 28 | 11 |
| 1/1:1/2 | 7 | 0 | 0 | 0 | 0 | 0 | 0 | 1 | 1 | 1 | 1 | 1 | 2 | 2 | 2 |
| 1/2:0/0 | 0 | 1 | 2 | 1 | 1 | 1 | 1 | 0 | 0 | 1 | 1 | 0 | 0 | 0 | 0 |
| 1/2:0/1 | 0 | 0 | 0 | 1 | 1 | 1 | 1 | 2 | 2 | 2 | 2 | 2 | 1 | 0 | 2 |
| 1/2:1/1 | 9 | 12 | 15 | 19 | 18 | 19 | 20 | 23 | 24 | 25 | 26 | 23 | 19 | 18 | 6 |
| concordant 0/0 | 14054746 | 19237587 | 21245357 | 22061259 | 22419936 | 22585939 | 22664220 | 22703546 | 22723853 | 22735585 | 22747418 | 22751474 | 22753357 | 22754464 | 22755575 |
| concordant 0/1 | 1845 | 2468 | 2936 | 3339 | 3711 | 4039 | 4308 | 4535 | 4754 | 4940 | 5261 | 5471 | 5616 | 5760 | 5964 |
| concordant 1/1 | 21578 | 29995 | 33441 | 35026 | 35754 | 36169 | 36402 | 36508 | 36612 | 36681 | 36801 | 36897 | 36981 | 37040 | 37159 |
| concordant 1/2 | 0 | 0 | 0 | 1 | 3 | 4 | 5 | 6 | 6 | 6 | 5 | 9 | 14 | 16 | 26 |
| discordant 0/1 | 1 | 0 | 0 | 2 | 2 | 1 | 1 | 1 | 1 | 1 | 2 | 4 | 3 | 2 | 0 |
| discordant 1/1 | 68 | 53 | 57 | 37 | 25 | 17 | 15 | 16 | 14 | 16 | 22 | 23 | 17 | 11 | 6 |

Table 12: Comparison of called genotypes in Vindija 33.19 subsamples to 30x calls. Genotypes column give the combination of genotypes in order 30x:subsample. Discordant genotypes list the events where the state matched, but the called alleles differed.

| Genotypes | 1x | 2x | 3x | 4x | 5x | 6x | 7x | 8x | 9x | 10x | 12.5x | 15x | 17.5x | 20x | 25x |
|---|---|---|---|---|---|---|---|---|---|---|---|---|---|---|---|
| 0/0:0/1 | 9 | 1 | 1 | 0 | 0 | 1 | 1 | 2 | 2 | 1 | 0 | 1 | 1 | 1 | 0 |
| 0/0:1/1 | 1515 | 1878 | 1299 | 806 | 463 | 290 | 169 | 114 | 73 | 40 | 13 | 3 | 0 | 0 | 0 |
| 0/1:0/0 | 266 | 774 | 735 | 628 | 528 | 411 | 332 | 270 | 246 | 204 | 134 | 87 | 52 | 29 | 5 |
| 0/1:1/1 | 1 | 0 | 0 | 0 | 0 | 0 | 0 | 0 | 0 | 0 | 0 | 0 | 0 | 0 | 0 |
| 1/1:0/0 | 8 | 7 | 4 | 5 | 3 | 0 | 0 | 0 | 0 | 0 | 0 | 0 | 0 | 0 | 0 |
| 1/2:0/1 | 0 | 0 | 0 | 0 | 0 | 0 | 0 | 1 | 1 | 0 | 0 | 0 | 0 | 0 | 0 |
| 1/2:1/1 | 0 | 3 | 6 | 5 | 4 | 3 | 2 | 3 | 3 | 2 | 1 | 1 | 1 | 1 | 0 |
| concordant 0/0 | 4298057 | 13965648 | 17863618 | 20021659 | 21188632 | 21822227 | 22167988 | 22364540 | 22483197 | 22557762 | 22653561 | 22695379 | 22717659 | 22730345 | 22742667 |
| concordant 0/1 | 217 | 694 | 767 | 916 | 1132 | 1447 | 1847 | 2225 | 2589 | 2946 | 3702 | 4254 | 4641 | 4941 | 5321 |
| concordant 1/1 | 5573 | 20989 | 27162 | 30751 | 32771 | 33982 | 34655 | 35098 | 35375 | 35587 | 35903 | 36096 | 36234 | 36313 | 36450 |
| concordant 1/2 | 0 | 0 | 0 | 1 | 1 | 2 | 2 | 2 | 3 | 4 | 4 | 5 | 9 | 10 | 10 |
| discordant 1/1 | 1 | 2 | 4 | 2 | 1 | 0 | 0 | 0 | 0 | 0 | 0 | 0 | 0 | 0 | 0 |

Table 13: Comparison of called genotypes of at least GQ30 in Vindija 33.19 subsamples to 30x calls. Labels as in Table 12.

| Genotypes | 1x | 2x | 3x | 4x | 5x | 6x | 7x | 8x | 9x | 10x | 12.5x | 15x | 17.5x | 20x | 25x |
|---|---|---|---|---|---|---|---|---|---|---|---|---|---|---|---|
| 0/0:0/1 | 0 | 0 | 0 | 0 | 0 | 0 | 0 | 0 | 0 | 0 | 0 | 1 | 0 | 0 | 0 |
| 0/1:0/0 | 0 | 19 | 42 | 50 | 60 | 47 | 56 | 49 | 44 | 34 | 26 | 18 | 6 | 2 | 0 |
| 1/2:1/1 | 0 | 0 | 0 | 0 | 0 | 1 | 1 | 1 | 1 | 1 | 0 | 0 | 0 | 0 | 0 |
| concordant 0/0 | 748 | 1241358 | 3509091 | 6215502 | 9257342 | 12043092 | 14401637 | 16302760 | 17780674 | 18908590 | 20692904 | 21590412 | 22064492 | 22322058 | 22565822 |
| concordant 0/1 | 4 | 31 | 89 | 222 | 398 | 617 | 949 | 1272 | 1615 | 2030 | 2900 | 3586 | 4107 | 4490 | 5011 |
| concordant 1/1 | 4 | 1394 | 4170 | 7831 | 12069 | 16264 | 19988 | 23158 | 25644 | 27635 | 30965 | 32820 | 33917 | 34557 | 35347 |
| concordant 1/2 | 0 | 0 | 0 | 0 | 0 | 1 | 1 | 2 | 2 | 2 | 4 | 5 | 5 | 5 | 6 |

Table 14: Comparison of called genotypes of at least GQ50 in Vindija 33.19 subsamples to 30x calls. Labels as in Table 12.

| Genotype | Vindija 33.15 without ref.bias | Vindija33.15 with ref.bias |
|---|---|---|
| AC | 0.0266134 | 0.0287228 |
| AG | 0.0963167 | 0.109077 |
| AT | 0.0244506 | 0.0271335 |
| CG | 0.0233174 | 0.0246703 |
| CT | 0.0962895 | 0.110025 |
| GT | 0.0257977 | 0.0272471 |

Table 15: Vindija 33.15: genotype frequency estimates with and without reference bias. The estimated reference bias for the third column was 10.14% ($r = 0.6014$). Genotype frequencies are given per 1000bp.

| GT Vi.33.15 | GT Vi.33.19 | ¬RB vs. ¬RB | RB vs. ¬RB | ¬RB vs. RB | RB vs. RB |
|---|---|---|---|---|---|
| 0/0 | 0/0 | 17461981 | 17461610 | 17461954 | 17461583 |
| 0/0 | 0/1 | 112 | 102 | 139 | 129 |
| 0/0 | 1/1 | 1 | 1 | 1 | 1 |
| 0/1 | 0/0 | 352 | 722 | 347 | 717 |
| 0/1 | 0/1 | 3785 | 3793 | 3790 | 3798 |
| 0/1 | 1/1 | 79 | 39 | 79 | 39 |
| 1/1 | 0/0 | 1 | 2 | 1 | 2 |
| 1/1 | 0/1 | 12 | 14 | 12 | 14 |
| 1/1 | 1/1 | 26117 | 26157 | 26117 | 26157 |
| 1/2 | 1/1 | 1 | 1 | 1 | 1 |
| 1/2 | 1/2 | 5 | 5 | 5 | 5 |

Table 16: Comparison of Vindija 33.19 and 33.15 chr21 genotypes with and without reference bias. The samples Vindija 33.19 and 33.15 have been found to originate from the same individual. The first two columns shows the observed configuration of genotypes. The remaining columns show the counts of genotypes of each configuration observed when calling Vindija 33.15 and Vindija 33.19 with (RB) or without (¬RB) taking reference bias into account. Parameter estimates for genotype frequencies with and without reference bias are listed in Tables 11 and 10, respectively, for Vindija 33.19 and in Table 15 for Vindija 33.15.

| Vi33.19 coverage | AC | AG | AT | CG | CT | GT |
|---|---|---|---|---|---|---|
| 0.9x | 0.0195129 | 0.109353 | 0.0205452 | 0.0187133 | 0.11283 | 0.022431 |
| 1.0x | 0.0193746 | 0.100528 | 0.0202986 | 0.0174671 | 0.101193 | 0.0219828 |
| 1.2x | 0.0191059 | 0.092621 | 0.0208953 | 0.0169481 | 0.0982858 | 0.0200446 |
| 1.5x | 0.0188508 | 0.0847961 | 0.0201072 | 0.016174 | 0.0860951 | 0.0185336 |
| 2.0x | 0.0189877 | 0.0808416 | 0.0205732 | 0.0161658 | 0.0862022 | 0.0179353 |
| 30x* | 0.0197930 | 0.065936 | 0.0205161 | 0.0195035 | 0.0646958 | 0.0203278 |

Table 17: Estimated genotype frequencies (per 1000bp) for subsampled Vindija 33.19 data and full data (30-fold coverage) for autosomal sites with at least 4-fold coverage. * 30x coverage estimate is the average, weighted by chromosome size, of the previously published genotype frequencies.

| Vi33.19 coverage | AC | AG | AT | CG | CT | GT |
|---|---|---|---|---|---|---|
| Goyet | 0.0167772 | 0.058848 | 0.0183608 | 0.0169625 | 0.0592227 | 0.0181386 |
| LesCottés | 0.0195414 | 0.0562096 | 0.0269036 | 0.014648 | 0.0600958 | 0.0209159 |
| Mez2 | 0.017562 | 0.0569594 | 0.0176347 | 0.0175739 | 0.0576068 | 0.0180015 |
| Vindija87 | 0.0201796 | 0.0758009 | 0.0240558 | 0.0203436 | 0.0809328 | 0.022701 |
| Spy | 0.031233 | 0.0893976 | 0.0394828 | 0.0264985 | 0.0949992 | 0.031182 |
| Loschbour 1x | 0.0510422 | 0.19892 | 0.0425446 | 0.0373921 | 0.195768 | 0.0535498 |
| Loschbour 2x | 0.0510101 | 0.201208 | 0.0409543 | 0.0388374 | 0.202462 | 0.052122 |
| Motala12 | 0.0549699 | 0.260431 | 0.0425594 | 0.0612579 | 0.259056 | 0.0556023 |
| Loschbour 22x* | 0.05067 | 0.20966 | 0.04319 | 0.05325 | 0.20831 | 0.05103 |

Table 18: Estimated genotype frequencies (per 1000bp) for low-coverage archaic and modern human data on autosomes for sites with at least 4-fold coverage. * Genome-wide average, weighted by chromosome sizes, of previous snpAD estimates.

| Parameter | No mapability | map35_100 |
|---|---|---|
| depth | 1.83309693 | 0.859994221 |
| fracMissing | 0.44624482 | 0.728850233 |
| fracTwoOrMore | 0.55375518 | 0.271149767 |
| pi(A) | 0.233486339 | 0.217336017 |
| pi(C) | 0.245584846 | 0.262575431 |
| pi(G) | 0.262368764 | 0.283677311 |
| pi(T) | 0.258560051 | 0.236411241 |
| theta_MLE | 0.00218729023 | 0.00123814321 |
| theta_C95_l | 0.00177227373 | 0.000745217841 |
| theta_C95_u | 0.00260230673 | 0.00173106857 |
| LL | -241726.674 | -117295.226 |

Table 19: ATLAS' genome-wide estimates of $\theta$ for Motala12 with and without mapability track.

### 3 Comparison to ATLAS

ATLAS 1.0 was downloaded on 2017-11-23 from the authors github page (rev. 9b2390c). Conserved regions were downloaded according to the authors instructions and modified to fit the coordinates in bam files:

```
cat original_hg19_UCNE_coord.bed |
        sed -e 's/^chr//g' > hg19_UCNE_coord.bed
```

ATLAS was then run with the *recal* option using the following steps:

```
atlas task=estimatePMD bam=motala12.bam \
        fasta=hg19_evan/whole_genome.fa \
        length=25

atlas task=recal bam=motala12.bam \
        pmdFile=motala12_PMD_input_Empiric.txt \
        regions=hg19_UCNE_coord.bed verbose

atlas task=recalBAM bam=motala12.bam \
        recal=motala12_recalibrationEM.txt \
        pmdFile=motala12_PMD_input_Empiric.txt \
        fasta=hg19_evan/whole_genome.fa \
        withPMD maxOutQuality=42 verbose
```

The resulting recalibrated quality scores appeared to shift all quality scores to values less than 10:

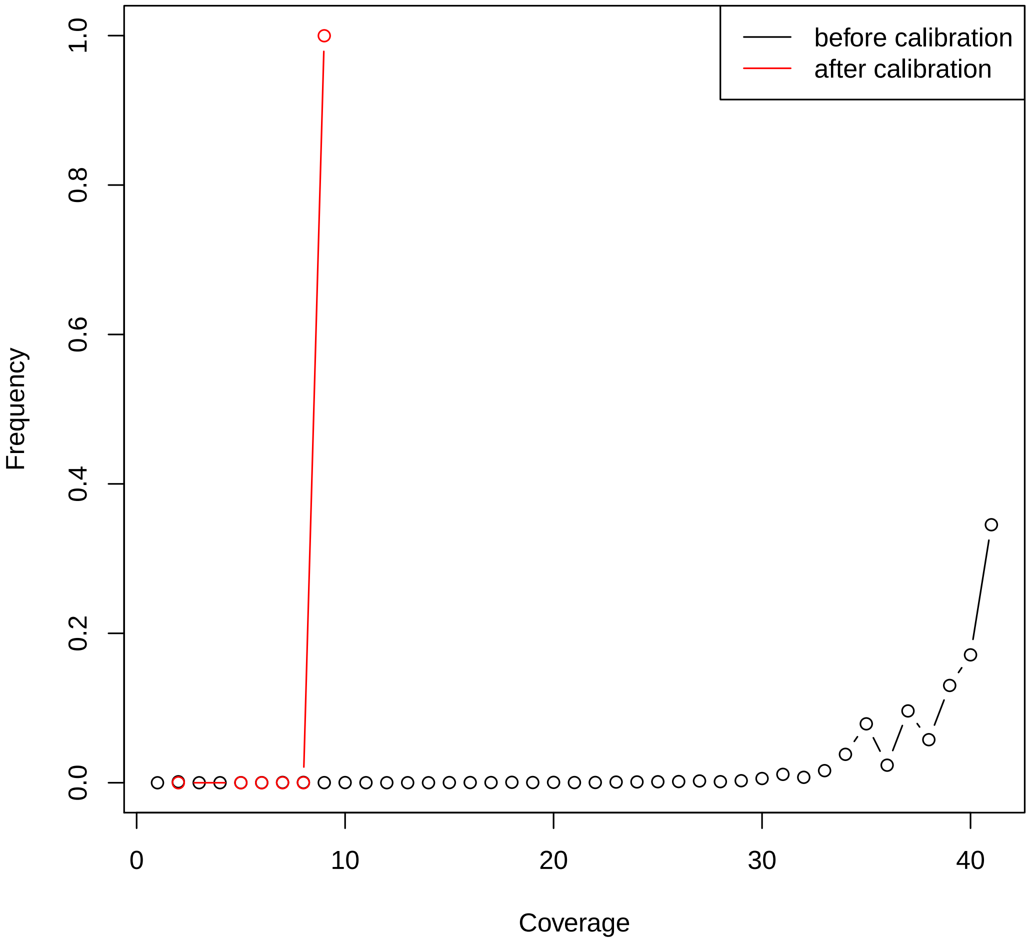

Running ATLAS with the *minDepth=2* option did not change the result.

A new version of ATLAS 1.0 was downloaded on 2018-03-15 (commit 49f1fea). PMD estimation was run with the command:

```
atlas task=estimatePMD \
        bam=motala12.bam \
        fasta=hg19_evan/whole_genome.fa \
        length=25
```

The command failed with the message "Error: Length mismatch!".

Using the previous estimates of the older ATLAS version for the *estimatePMD* step, the following commands were run:

```
atlas task=recal bam=motala12.bam \
        pmdFile=motala12_PMD_input_Empiric.txt \
        regions=hg19_UCNE_coord.bed verbose

atlas task=recalBAM bam=motala12.bam \
        recal=motala12_recalibrationEM.txt \
        pmdFile=motala12_PMD_input_Empiric.txt \
        fasta=hg19_evan/whole_genome.fa withPMD \
        maxOutQuality=42 verbose
```

The execution of *recalBAM* failed with the same message as before, but left an indexable recalibrated bam file with a more reasonable quality score distribution:

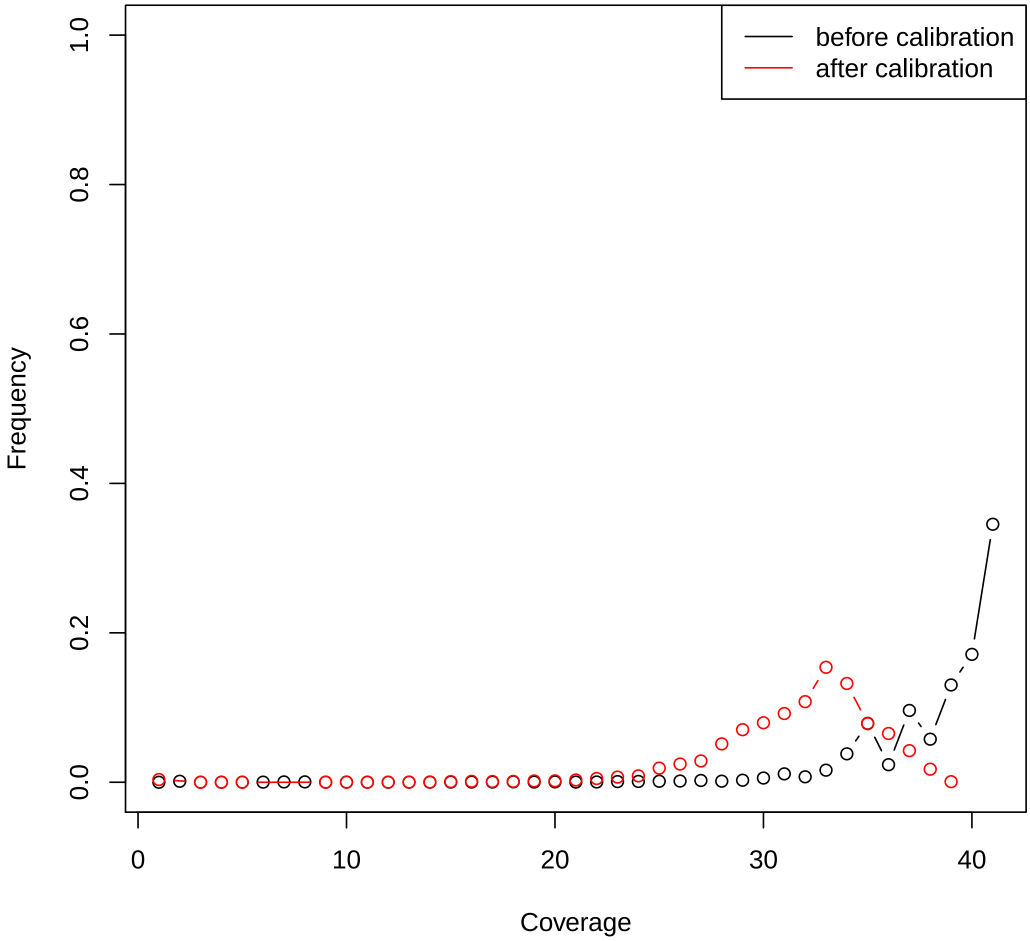

Genome-wide theta on the autosomes were estimated for all regions and for regions passing a 35mer mapability filter (map35_100) that was used in the analysis of the Altai Neandertal and Vindija Neandertal, and for all snpAD estimates in this paper:

```
atlas task=estimateTheta bam=motala12.bam \
        pmdFile=motala12_PMD_input_Empiric.txt \
        recal=motala12_recalibrationEM.txt \
        thetaGenomeWide limitChr=22 \
        minDepth=2 verbose

atlas task=estimateTheta bam=motala12.bam \
        pmdFile=motala12_PMD_input_Empiric.txt \
        recal=motala12_recalibrationEM.txt \
        thetaGenomeWide limitChr=22 \
        regions=hs37m_filt35_99.bed.gz \
        minDepth=2 verbose
```

The resulting estimates are shown in Table 19.

Lazaridis, I., Patterson, N., Mittnik, A., Renaud, G., Mallick, S., Kirsanow, K., Sudmant, P. H., Schraiber, J. G., Castellano, S., Lipson, M., Berger, B., Economou, C., Bollongino, R., Fu, Q., Bos, K. I., Nordenfelt, S., Li, H., de Filippo, C., Prüfer, K., Sawyer, S., Posth, C., Haak, W., Hallgren, F., Fornander, E., Rohland, N., Delsate, D., Francken, M., Guinet, J.-M., Wahl, J., Ayodo, G., Babiker, H. A., Bailliet, G., Balanovska, E., Balanovsky, O., Barrantes, R., Bedoya, G., Ben-Ami, H., Bene, J., Berrada, F., Bravi, C. M., Brisighelli, F., Busby, G. B. J., Cali, F., Churnosov, M., Cole, D. E. C., Corach, D., Damba, L., van Driem, G., Dryomov, S., Dugoujon, J.-M., Fedorova, S. A., Gallego Romero, I., Gubina, M., Hammer, M., Henn, B. M., Hervig, T., Hodoglugil, U., Jha, A. R., Karachanak-Yankova, S., Khusainova, R., Khusnutdinova, E., Kittles, R., Kivisild, T., Klitz, W., Kučinskas, V., Kushniarevich, A., Laredj, L., Litvinov, S., Loukidis, T., Mahley, R. W., Melegh, B., Metspalu, E., Molina, J., Mountain, J., Näkkäläjärvi, K., Nesheva, D., Nyambo, T., Osipova, L., Parik, J., Platonov, F., Posukh, O., Romano, V., Rothhammer, F., Rudan, I., Ruizbakiev, R., Sahakyan, H., Sajantila, A., Salas, A., Starikovskaya, E. B., Tarekegn, A., Toncheva, D., Turdikulova, S., Uktveryte, I., Utevska, O., Vasquez, R., Villena, M., Voevoda, M., Winkler, C. A., Yepiskoposyan, L., Zalloua, P., Zemunik, T., Cooper, A., Capelli, C., Thomas, M. G., Ruiz-Linares, A., Tishkoff, S. A., Singh, L., Thangaraj, K., Villems, R., Comas, D., Sukernik, R., Metspalu, M., Meyer, M., Eichler, E. E., Burger, J., Slatkin, M., Pääbo, S., Kelso, J., Reich, D., and Krause, J. (2014). Ancient human genomes suggest three ancestral populations for present-day Europeans. *Nature*, **513**(7518), 409–413.

Prüfer, K., Racimo, F., Patterson, N., Jay, F., Sankararaman, S., Sawyer, S., Heinze, A., Renaud, G., Sudmant, P. H., de Filippo, C., Li, H., Mallick, S., Dannemann, M., Fu, Q., Kircher, M., Kuhlwilm, M., Lachmann, M., Meyer, M., Ongyerth, M., Siebauer, M., Theunert, C., Tandon, A., Moorjani, P., Pickrell, J., Mullikin, J. C., Vohr, S. H., Green, R. E., Hellmann, I., Johnson, P. L. F., Blanche, H., Cann, H., Kitzman, J. O., Shendure, J., Eichler, E. E., Lein, E. S., Bakken, T. E., Golovanova, L. V., Doronichev, V. B., Shunkov, M. V., Derevianko, A. P., Viola, B., Slatkin, M., Reich, D., Kelso, J., and Pääbo, S. (2014). The complete genome sequence of a Neandertal from the Altai Mountains. *Nature*, **505**(7481), 43–49.

Prüfer, K., de Filippo, C., Grote, S., Mafessoni, F., Korlević, P., Hajdinjak, M., Vernot, B., Skov, L., Hsieh, P., Peyrégne, S., Reher, D., Hopfe, C., Nagel, S., Maricic, T., Fu, Q., Theunert, C., Rogers, R., Skoglund, P., Chintalapati, M., Dannemann, M., Nelson, B. J., Key, F. M., Rudan, P., Kućan, Ž., Gušić, I., Golovanova, L. V., Doronichev, V. B., Patterson, N., Reich, D., Eichler, E. E., Slatkin, M., Schierup, M. H., Andrés, A. M., Kelso, J., Meyer, M., and Pääbo, S. (2017). A high-coverage Neandertal genome from Vindija Cave in Croatia. *Science (New York, N.Y.)*, **358**(6363), 655–658.
